# Differentially Expressed Heterogeneous Overdispersion Genes Testing for Count Data*

**DOI:** 10.1101/2023.02.21.529455

**Authors:** Yubai Yuan, Qi Xu, Agaz Wani, Jan Dahrendor, Chengqi Wang, Janelle Donglasan, Sarah Burgan, Zachary Graham, Monica Uddin, Derek Wildman, Annie Qu

## Abstract

The mRNA-seq data analysis is a powerful technology for inferring information from biological systems of interest. Specifically, the sequenced RNA fragments are aligned with genomic reference sequences, and we count the number of sequence fragments corresponding to each gene for each condition. A gene is identified as differentially expressed (DE) if the difference in its count numbers between conditions is statistically significant. Several statistical analysis methods have been developed to detect DE genes based on RNA-seq data. However, the existing methods could suffer decreasing power to identify DE genes arising from overdispersion and limited sample size. We propose a new differential expression analysis procedure: heterogeneous overdispersion genes testing (DEHOGT) based on heterogeneous overdispersion modeling and a post-hoc inference procedure. DEHOGT integrates sample information from all conditions and provides a more flexible and adaptive overdispersion modeling for the RNA-seq read count. DEHOGT adopts a gene-wise estimation scheme to enhance the detection power of differentially expressed genes. DEHOGT is tested on the synthetic RNA-seq read count data and outperforms two popular existing methods, DESeq and EdgeR, in detecting DE genes. We apply the proposed method to a test dataset using RNAseq data from microglial cells. DEHOGT tends to detect more differently expressed genes potentially related to microglial cells under different stress hormones treatments.

## 1 Introduction

High-throughput sequencing of DNA fragments and mRNA-seq techniques are powerful tools based on next generation sequencing technologies [16] for monitoring RNA abundance to detect genetic variation. Specifically, for RNAseq, the sequenced RNA fragments are aligned with reference genome sequences, and the number of sequence fragments assigned to each gene is counted for each sample. Then we can compare read counts between different biological conditions or between different genetic variants to infer genetic information based on biological systems of interest [19]. In the analysis of RNA-seq data, read counts do not have a prior upper bound, thus regression models based on a binomial distribution with a pre-specified number of trails do not apply [34]. Linear regression is therefore not feasible as count data is always a non-negative integer. More importantly, RNA-seq data presents high overdispersion, implying that the variance of the count can be much larger than its mean. Given that the sample sizes are typically small for RNA-seq analysis due to the cost and other factors, statistical modeling needs to address the large variation from the data and to improve the power of detecting differential gene expressions.

One fundamental clinical interest of applying RNA-seq analysis is to understand the mechanism of post-traumatic stress disorder (PTSD) formulation. PTSD is a common severe psychiatric disorder that develops following exposure to a life-threatening or traumatic experience [31]. PTSD is known to cause negative effect on an individual’s life quality via the PTSD condition itself or the relevant comorbidities. Previous works [12, 18] show that only a small proportion of individuals experience traumatic events will develop PTSD. Meanwhile, the majority of people exposed to trauma are resilient even after repeated exposures to trauma [35]. In addition, various risk factors of PTSD have been identified such as low socio-economic status, social support and gender [33, 6, 15].

Significant individual heterogeneity of either response to trauma or the PTSD development originates from the individual epigenetic variability. Specifically, previous studies reveal the connection between PTSD and immune system functioning, and several genes such as FKBP5 involved with the immune system are also found to be differentially expressed among PTSD individ-uals [27, 32, 17]. In particular, previous work [13] has identified monocytes as a key cell type in differentiating male subjects with versus without lifetime PTSD. In addition, rodent studies have implicated peripheral monocytes in inducing anxiety-like behavior through trafficking of proinflammatory monocytes to the brain via activated microglia. Following this line of research, in this paper, we collect RNA-seq data from the well-designed lab experiments to investigate differential expression of genes in human microglia cells under different immune characteristic environments. This is an important step for understanding the role of microglia cells and immune-related genes in PTSD development.

The main challenge in analyzing microglial RNA-seq datasets lies in the high and heterogeneous overdispersion in the read counts. As an illustration, Figure 1 shows the histogram of the empirical RNA read counts from microglial data, where the read counts are highly spread out and the variance can be much larger than the mean. Several differential expression analysis methods have been developed to address the overdispersion issue in RNA-seq read counts. Among these methods, the DESeq [1] and EdgeR [23] are the most popular and are implemented and available using the R [8]. Specifically, the DESeq analyzes count data by using a shrinkage estimation for dispersions as well as fold changes to improve stability and interpretability of estimates. EdgeR is designed for the analysis of replicated count-based expression data, and is based on the method developed by Robinson and Smyth [25] using an overdispersed Poisson model to account for the read count variability. However, most existing methods adopt the shrinkage strategy when estimating the level of overdispersion by assuming that genes with similar expression strength have homogeneous dispersion levels. Although overdispersion regularization helps to increase the robustness of inference against the uncertainty due to limited sample size, it decreases the discriminative power in detecting differentially expressed genes with strong overdispersion effects at the population level.

**Figure 1:**
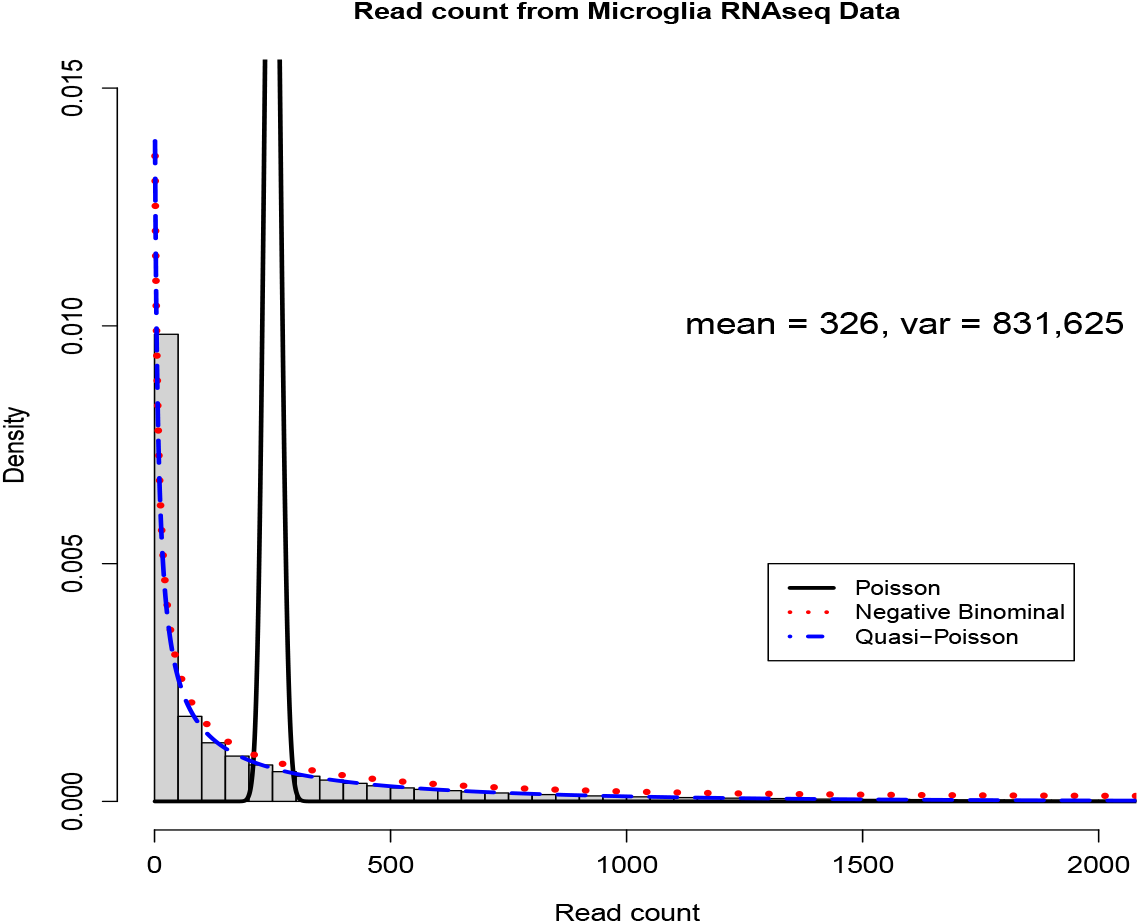
The overdispersion in the real RNA-seq count data

In this paper, we propose a new differential expression analysis framework based on generalized linear modeling. Compared with other popular RNA-seq analysis methods such as DESeq and EdgeR, the main advantages of the proposed method for differentially expressed heterogeneous overdispersion genes testing (DEHOGT) are as follows. First, our method jointly estimates the fold change and overdispersion parameters over samples from all treatment conditions, which increases the effective sample size and leads to more accurate inference. Second, and more importantly, our model adopts a within-sample independent structure among genes without assuming that genes with similar expression strength have homogeneous dispersion levels. Therefore, our method can better account for the heterogeneity in count dispersion and select more relevant genes. Third, our method allows for fully independent gene-wise inference and hence can achieve computational scalability to handle large gene datasets by implementing parallel computing. Finally, the proposed method enjoys the flexibility of adapting different overdispersion patterns by allowing different count generating distributions in the inference procedure.

## 2 Methodology

We develop a new differentially expressed gene testing procedure to account for the heterogeneity in gene-wise overdispersion levels. Traditionally, Poisson and multinomial distributions are used to model count data with large variance. However, the variance of RNA sequence counts tends to be much larger than that of the Poisson or multinominal distribution [30]. Overlooking the overdispersion could result in biased and misleading inference about gene association to the response of interest. To overcome this limitation, we first introduce adaptive distribution modeling in this paper to analyze the overdispersed RNA-seq count data. We utilize a quasi-Poisson distribution and a negative binominal distribution as the read count, thus generating a distribution similar to the overdispersion pattern which is based on empirical data. Specifically, we denote *Y* as the random count response, and the quasi-Poisson distribution satisfies:

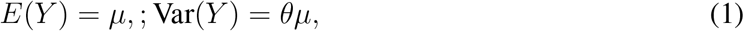

where *μ* > 0 is the mean of *Y, θ* ≥ 1 denotes the overdispersion parameter, and larger *θ* indicates higher overdispersion level. Although *μ* is larger than 0, *Y* can be any nonnegative integer. Note that Poisson model assumes that the variance is equal to the mean, e.g., *θ* = 1. In contrast, a quasi-Poisson distribution provides more flexibility to allow variance increases as a linear function of the mean. Accordingly, the quasi-Poisson regression generalizes the Poisson regression and is adopted to model an overdispersed count variable. The quasi-Poisson model is characterized by the first two moments, i.e., mean and variance. Besides the quasi-Poisson distribution, the negative binominal distribution can also be used to model overdispersed count data satisfying:

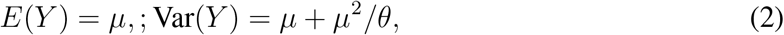

where *θ* > 0 is the overdispersion parameter, and smaller *θ* indicates higher overdispersion level. Similar to the quasi-Poisson distribution, the negative binominal distribution is characterized by the mean and variance while modeling the variance as a quadratic function of the mean. In the following, we denote the quasi-Poisson distribution and negative distribution as quasi-Poisson(*μ, θ^QP^*) and negative-binomial(*μ, θ^NB^*), respectively. In addition, we use *NB* and *QP* as the abbreviation of negative binomial and quasi-Poisson distribution. In Figure 1, we illustrate the distribution density functions by fitting the empirical read counts in our empirical data from microglia cells (see Methods) with 1) Poisson distribution, 2) negative binomial distribution, and 3) quasi-Poisson distribution, respectively. Compared with the Poisson distribution, both the negative binomial and quasi-Poisson distributions provide better approximation by capturing the overdispersion in read counts.

In addition, the read counts of a gene can be affected by other factors in an experiment other than its expression level in the RNA-seq. Therefore, instead of directly modeling the raw count data *Y*, we first perform count normalization, which makes the expression levels of genes more comparable and accurate between samples. We utilize the Trimmed Mean of M-values normalization (TMM) [24] adopted by EdgeR to compute the normalization factors that correct sample-specific biases. TMM is recommended for most RNA-Seq data where most genes are not differentially expressed across any pairs of the samples. Specifically, we first calculate the normalization factors as the median ratio of gene counts relative to the geometric mean per gene within a specific sample. The normalization factors account for two main non-expression factors; e.g., sequencing depth and RNA composition before between-sample comparison [24]. Consequently, we divide raw counts by sample-specific size factors to yield the effective read count for cross-sample comparisons.

The proposed DEHOGT workflow combines the above ingredients to identify differentially expressed genes. Compared with the two popular RNA-seq analysis methods DESeq and EdgeR, the main difference of the proposed method is at the model fitting step of the above algorithm, where the overdispersion parameters {*θ_i_*} are estimated for each gene individually. The DESeq and EdgeR estimate the overdispersion parameters by pooling the samples from different genes under the assumption that genes with similar expression strength also share similar overdispersion levels. In contrast, the proposed method does not rely on the homogeneous dispersion assumption and can capture the heterogeneity in different genes’ expression levels, especially when the overdispersion of gene is high. In addition, the proposed method allows one to choose different working distributions in Step 3 to model the RNA-seq count data to accommodate different associations between mean and variance presented in the empirical read count data. This provides us additional flexibility in modeling the overdispersion patterns to achieve more accurate read count fitting. Consequently, correctly specified read count overdispersion patterns can lead to higher statistical power of post-hoc testing to detect differentially expressed genes.

We summarize the proposed method (DEHOGT) for the RNAseq read count for detecting differentially expressed (DE) genes as follows. Assume that there exists a total of *R* different treatments and *S* samples where each treatment has multiple samples as replicated measurements. We index the gene and sample measurements as *g* and *s* such that *g* = 1,2, ⋯, *N* and *s* = 1, 2, ⋯, *S*. First, the read count data is modeled via one of the following generating distributions:

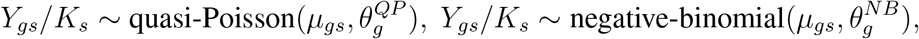

where *K_s_* denotes the normalization factor for the *s*th sample obtained by the TMM method. To determine the generating distribution, we check the overdispersion pattern between *E_s_*(*Y_gs_/K_s_*) and Var_*s*_ (*Y_gs_/K_s_*) from the empirical data. A better quadratic function fitting leads to the choice of a negative binomial distribution and a better linear relation fitting leads to the quasi-Poisson. Here we assume that the gene-wise dispersion level is constant across all samples to estimate the quasiPoisson distribution *θ^QP^*, or the negative binominal distribution *θ^NB^*, by utilizing information from samples under different treatments.

To differentiate genes’ read counts under different treatments, we model the genewise read count mean via the following generalized linear model:

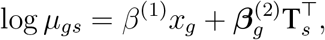

where 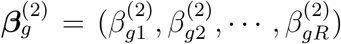 represents the fold change of the *g*th gene under *R* different treatments, and **T**_*s*_ ∈ {0,1}^*R*^ is the dummy coding for the treatment membership of the sth sample such that **T**_*sr*_ = 1 when the sth sample belongs to treatment *r, r* = 1, ⋯, *R*. In addition, *x_g_* ∈ ℝ^*p*^ are the gene-wise covariates, so that our method can further adjust other non-expression factors to reduce the bias in inferring the genes’ expression level, where *R* denotes a real number.

Given that 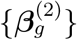 represents the gene-wise expression level under different treatments, we can infer whether the *g*th gene is differentially expressed under the two treatments *r*_1_ and *r*_2_ based on the linear hypothesis testing 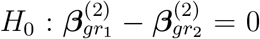. Then we can identify the DE genes under the treatment comparison pair (*r*_1_, *r*_2_) when the corresponding p-value is smaller than a specific cutoff. To control the type-I error of simultaneously testing on multiple genes, we adopt the Benjamini-Hochberg procedure [3] to adjust the gene-wise p-value, and control the false discovery rate. In addition to the adjusted p-value, the magnitude of the logfold change is also suggested as another criterion for choosing DE genes with a logfold change of 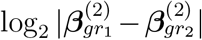 larger than 1.5 [20, 21]. Therefore, we combine these two criteria, and select the DE genes with an adjusted p-value smaller than 0.05 and an absolute logfold change larger than 1.5.

Our proposed DEHOGT algorithm is summarized as follows:

### Algorithm 1: DEHOGT

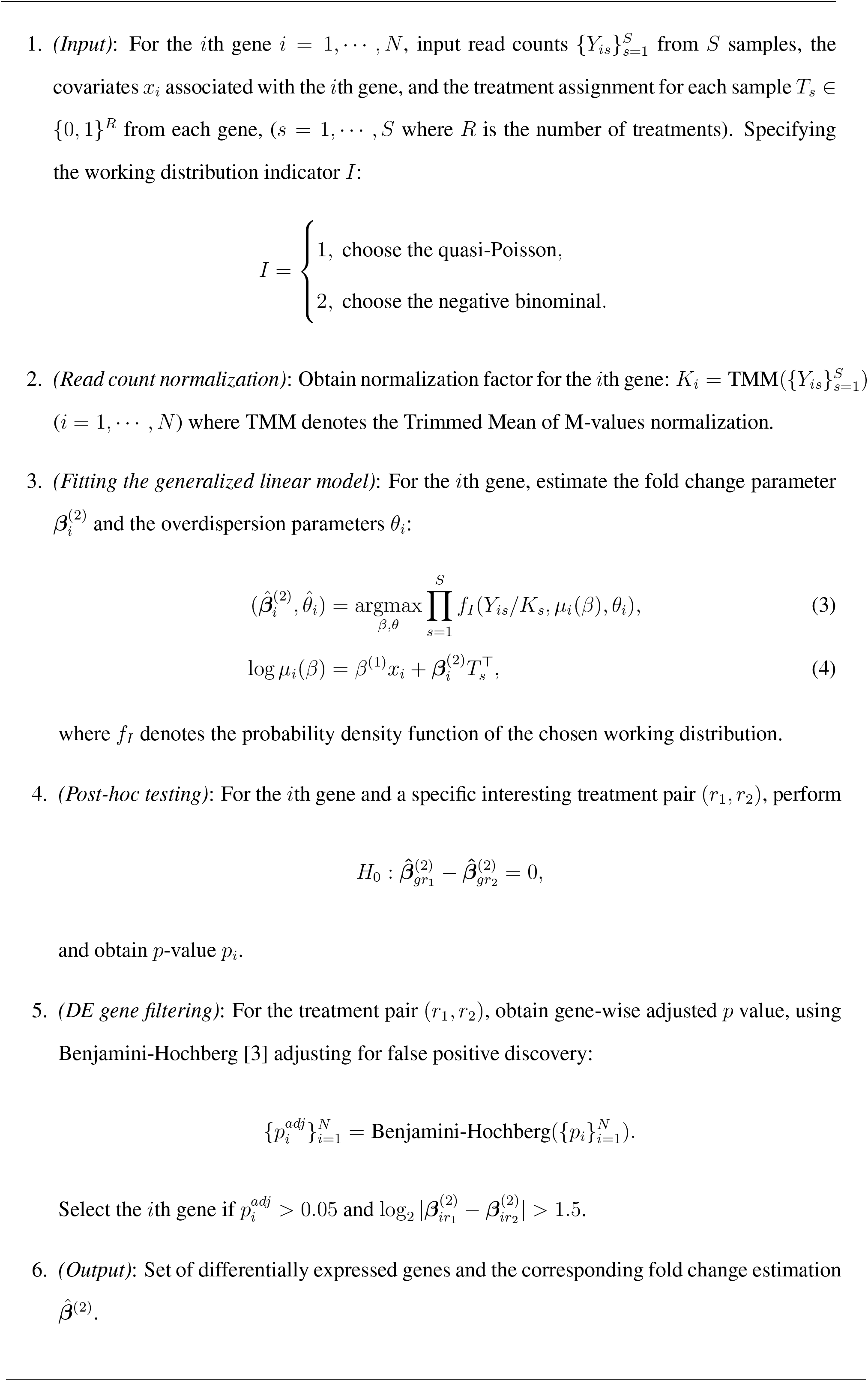

## 3 Simulation Studies

We compare the proposed DEHOGT method with two popular RNA-seq analysis methods DESeq [1] and EdgeR [23] in detecting differentially expressed genes on the simulated read count data and microglia cell RNA-seq data. In the first simulation setting, the discrepancy in expression level between the treatment and control group is weak for DE genes, while the average expression levels for both groups are high. In the second simulation setting, the expression discrepancy between the treatment and control group is strong for DE genes, while the average expression levels for both groups are low.

### 3.1 Read count with low discrepancy of expression level

In the first setting, we simulate the read count data following the negative binomial and the quasi-Poisson distribution:

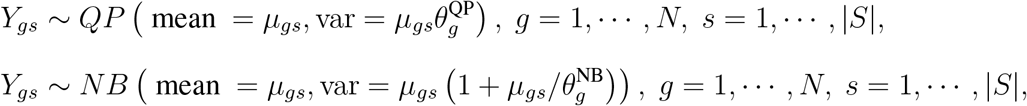

where *g* ∈ {1, ⋯, *N*} denotes gene indexes and the total number of genes *N* = 12,500. We use *G^DE^* ⊂ {1, ⋯, *N*} to denote the set of differentially expressed genes with |*G^DE^*| = 2500. In addition, *s* ∈ *S* and |*S*| = 12 denote the sample index with *S* = *S*_1_ ∪ *S*_2_, |*S*_1_| = |*S*_2_| = 6, where *S*_1_ and *S*_2_ indicate the samples in the control group and the treatment group, respectively. Here the mean parameters *μ_gs_* are similar to setting [29] in the RNA-seq data analysis. Specifically, the formulations are:

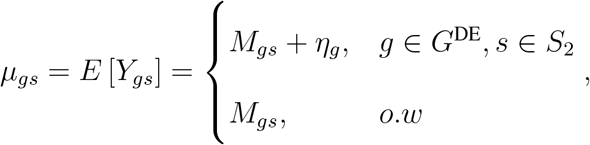

where we sample *M_gs_* from Unif[0, *U_s_*], and *U_s_* ~ Unif[600, 800] is the sample-wise sequencing depth. Furthermore, we sample *η_g_* from exp(1/100) as the up-regulated signal of the differentially expressed genes. We consider three different overdispersion levels for the read counts from the quasi-Poisson distribution as

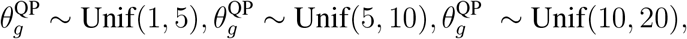

where a larger 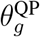 indicates a greater overdispersion level. Similarly, we consider three overdispersion levels under the negative binomial read counts as

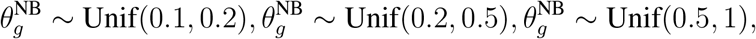

where a smaller 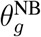 indicates a greater level of overdispersion.

We compare the performance of DESeq [1], EdgeR [23], and the proposed DEHOGT in identifying differentially expressed genes using an adjusted p value less than 0.05 and an absolute value of logfold change larger than 1.5. We first investigate the false negative rates from the comparison methods. The results under different data generations (quasi-Poisson or negative binomial) and different overdispersion levels are shown in Figures 2 and 3. Figures 2 and 3 suggest that the proposed DEHOGT method reaches the lowest false negative rate over competing methods under different overdispersion levels, indicating that most of the genes selected by the proposed method are differentially expressed. Note that the DEHOGT (NB) under the true negative binominal setting always achieves the lowest false negative rate when the cutoff of the adjusted p-value is set as 0.05. This is because the p-values from DEHOGT under NB tend to be smaller than for DEHOGT under QP. The better performance of DEHOGT under QP for the ROC and AUC (area under the ROC curve) implies that we can select a p-value cutoff larger than 0.05, under which the false negative rate of DEHOGT under QP can be smaller than the false negative rate of DEHOGT under NB.

**Figure 2:**
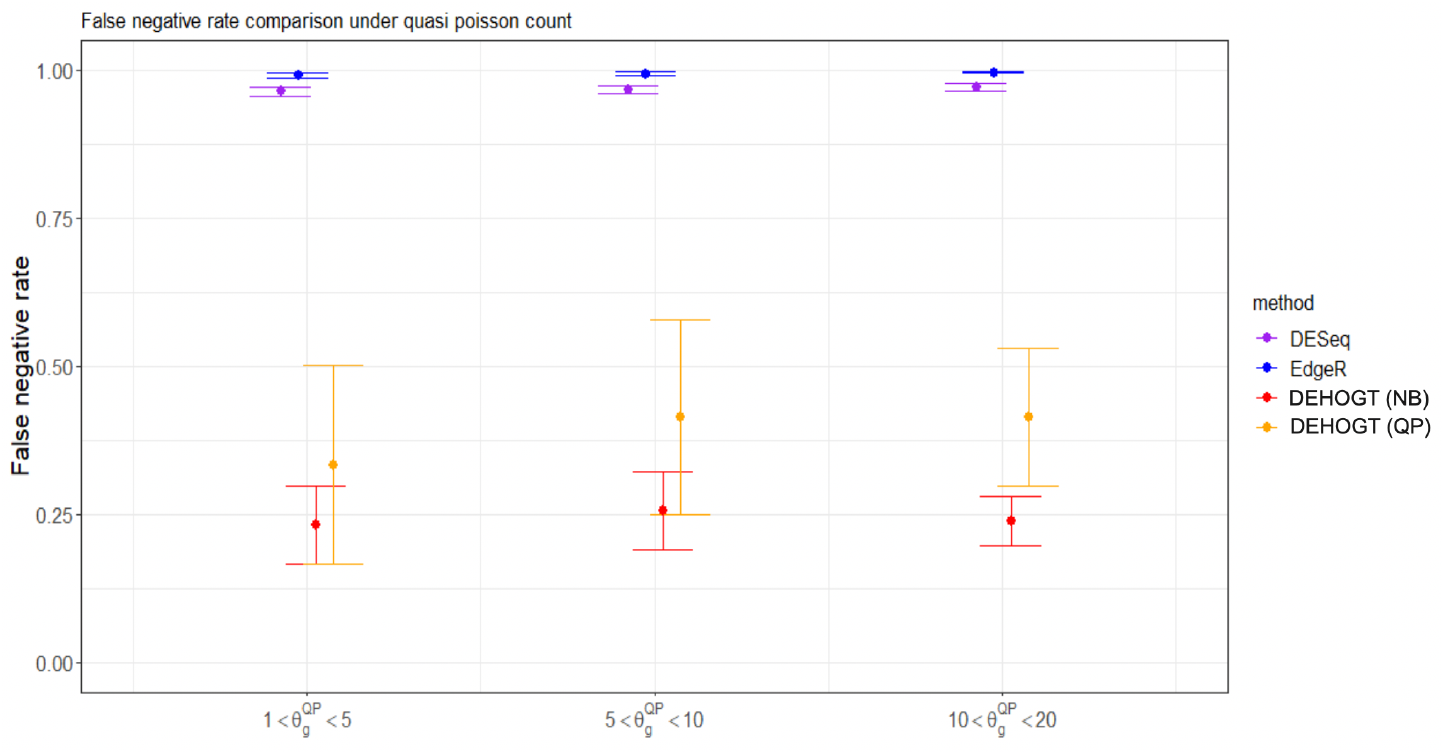
The false negative rate from different methods when the read counts follow the quasi-Poisson distribution with different overdispersion levels 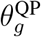 in simulation setting 1. The bars represents the standard deviation of the false negative rate over repeated experiments.

**Figure 3:**
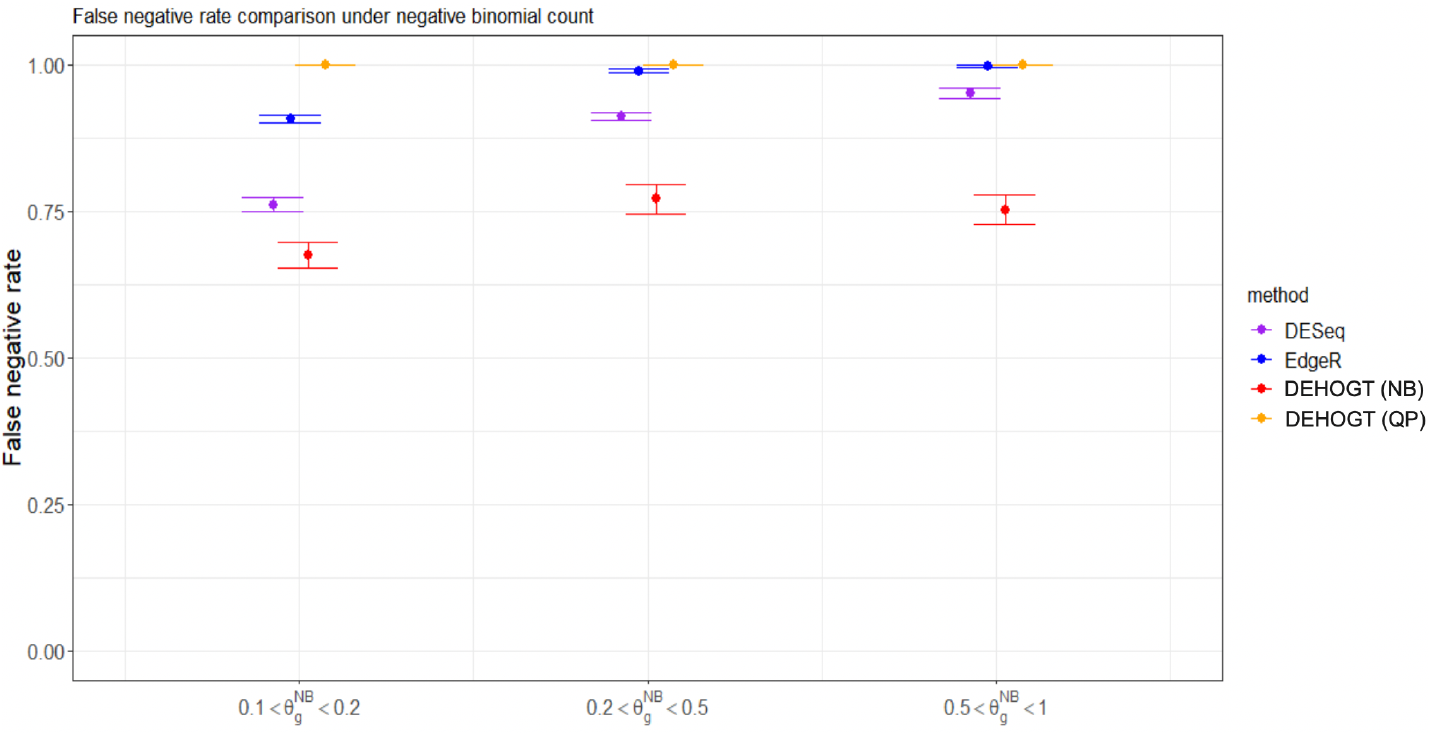
The false negative rate from different methods when the read counts follow the negative binomial distribution with different overdispersion levels 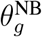 in simulation setting 1. The bars represents the standard deviation of the false negative rate over repeated experiments.

We also investigate the overall DE gene discriminative power of different methods when the cutoff point of the adjusted p-value changes over the range from 0 to 1, as measured by the AUC (area under the ROC curve). Note that the AUC value is between 0 and 1, and a larger AUC value indicates that the algorithm can achieve an overall lower false positive rate and lower false negative rate simultaneously. The comparisons are shown in Figures 4 and 5, illustrating the AUC values for competing methods under different generating distributions and overdispersion levels.

**Figure 4:**
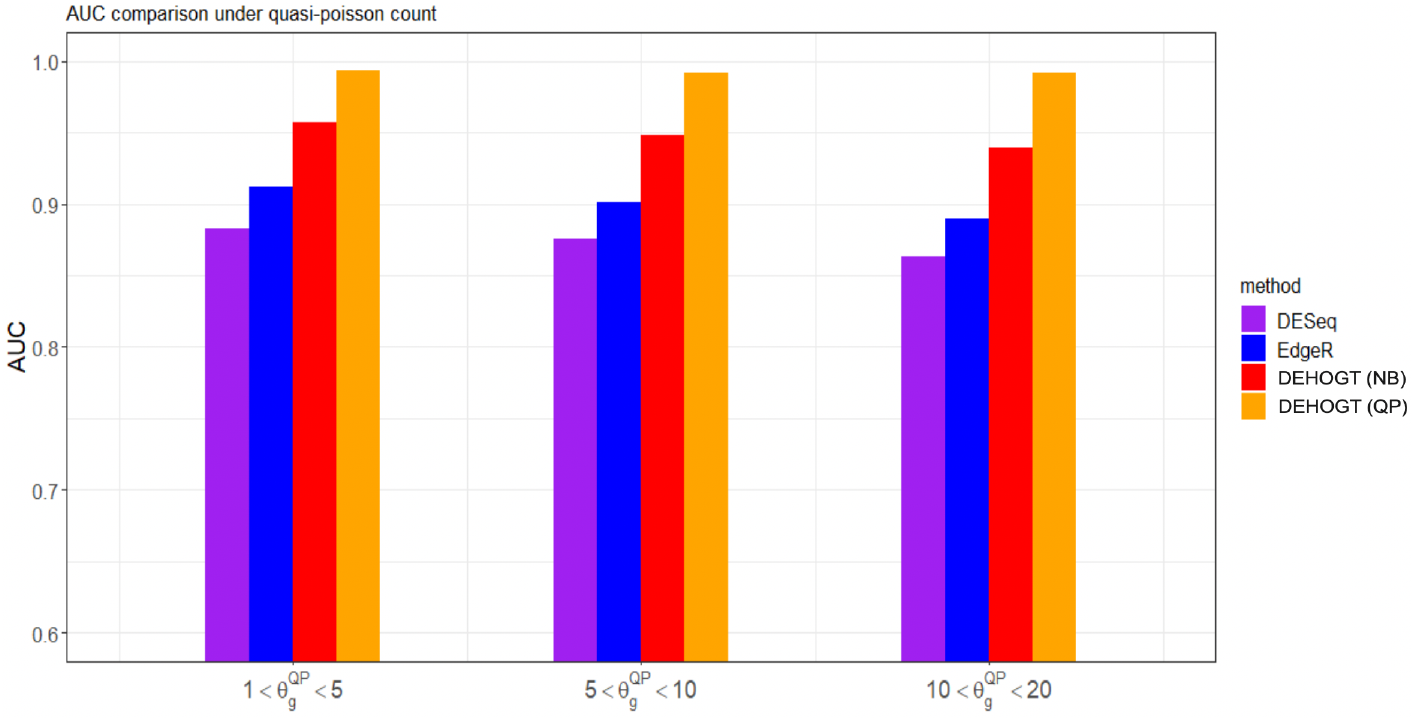
The area under the ROC curve from different methods when the read counts follow the quasi-Poisson distribution with different overdispersion levels 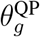 in simulation setting 1.

**Figure 5:**
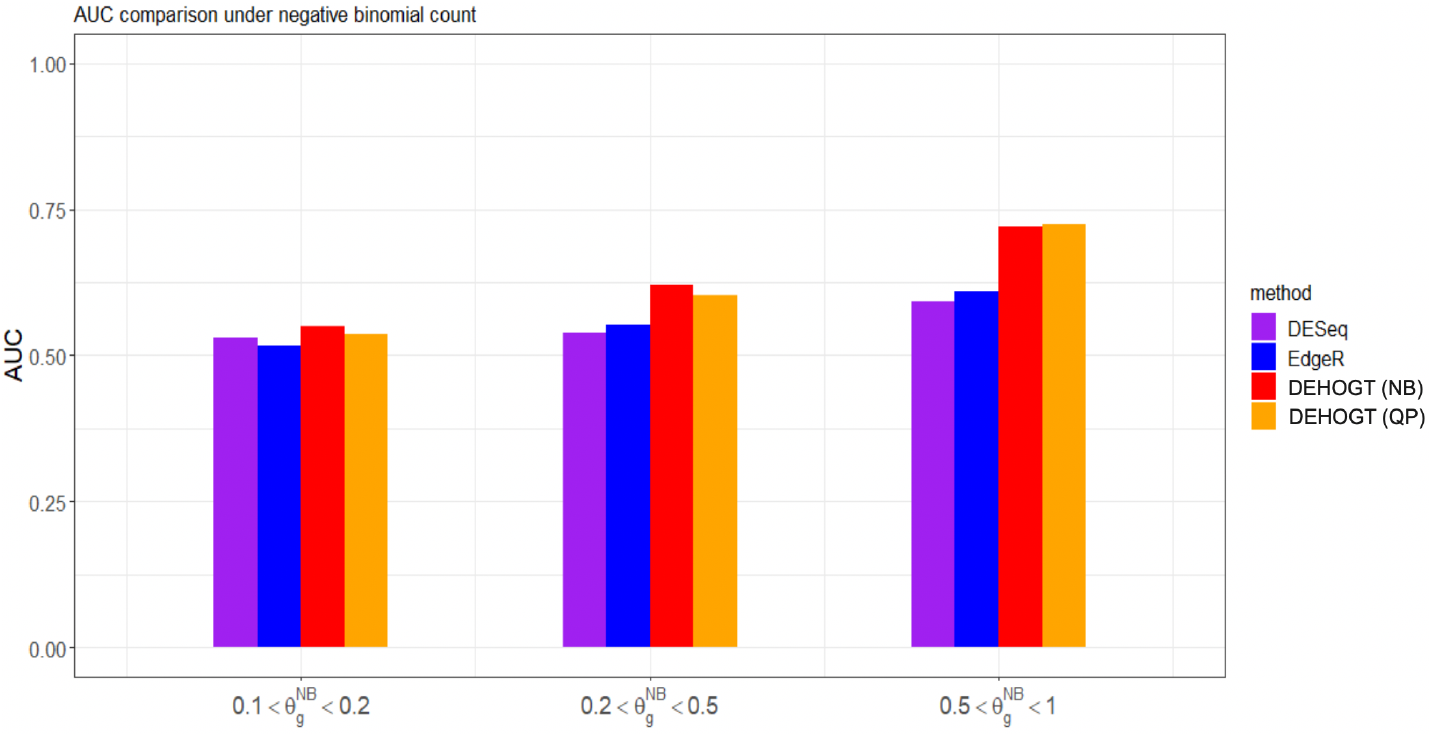
The area under the ROC curve from different methods when the read counts follow the negative binomial distribution with different overdispersion levels θNB in simulation setting 1.

The above results indicate that the proposed DEHOGT method outperforms both the DESeq and edgeR methods, and the proposed method can achieve the optimal AUC if the model is correctly specified. Specifically, DEHOGT under QP attains a higher AUC than DEHOGT under NB under varying 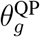 when the read counts are generated from the quasi-Poisson distribution. Similarly, DEHOGT (NB) attains higher AUC then DEHOGT (QP) under varying 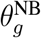 if the read counts are generated from negative binomial distributions.

### 3.2 Read count with high discrepancy of expression level

In the second simulation setting, we simulate the read count data of the moderate overdispersion level in RNAseq read counts. Following the notations in simulation 1, we simulate the read count data from both the quasi-Poisson distribution and the negative binomial distribution as

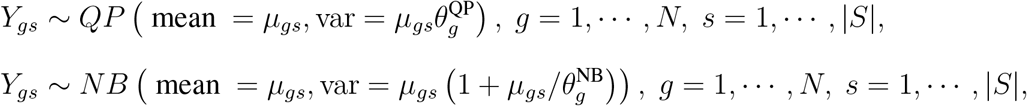

where we choose *N* = 10, 000 and *S* = *S*_1_ ∪ *S*_2_, |*S*_1_| = |*S*_2_| = 6. The GE genes are randomly selected and |*G^DE^*| = 2,000. We consider three different overdispersion levels for the read counts from the quasi-Poisson distribution as

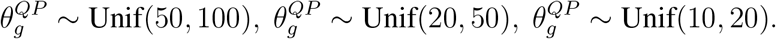

Similarly, we also consider three overdispersion levels under negative binomial read counts as

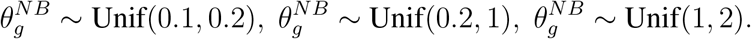

We differentiate DE genes and non-DE genes with different sample means such that

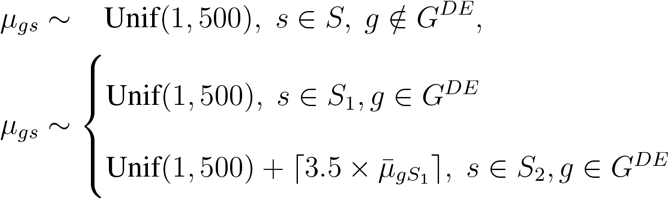

where ⌈·⌉ is the ceiling function, and 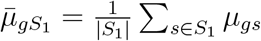.

Notice that the expression discrepancy between the treatment and control group is strong for DE genes, while the average expression levels for both groups are low. To select the DE genes, we follow the selection criterion in the previous simulation such that the absolute value of log2fold change is larger than 1.5 and the adjusted p value is smaller than 0.05. We first investigate the false negative rates from different methods, and the results are illustrated in Figure 6 and 7.

**Figure 6:**
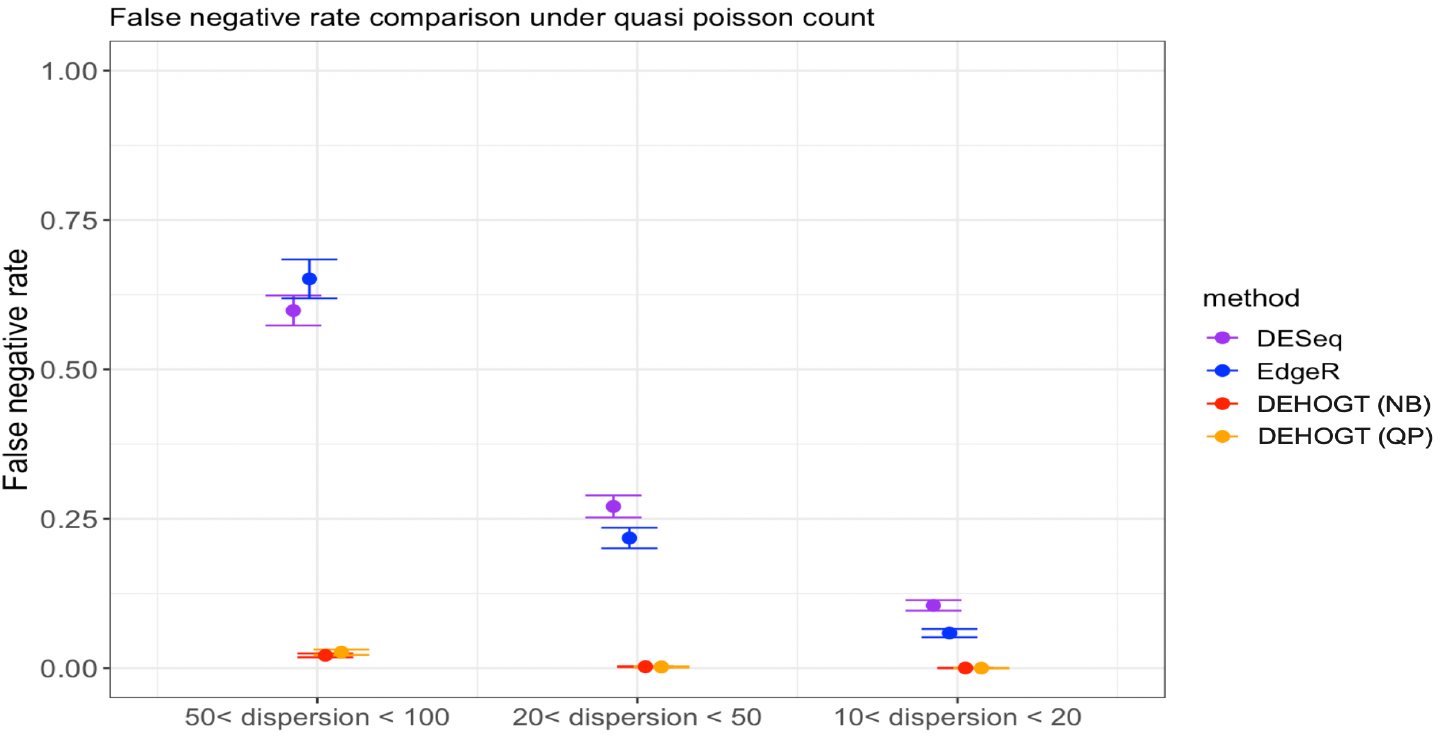
The false negative rate from different methods when the read count follow the quasi-Poisson distribution with different overdispersion levels *θ^QP^*. The variance of FNR obtained from repeated experiments is illustrated using the bars.

**Figure 7:**
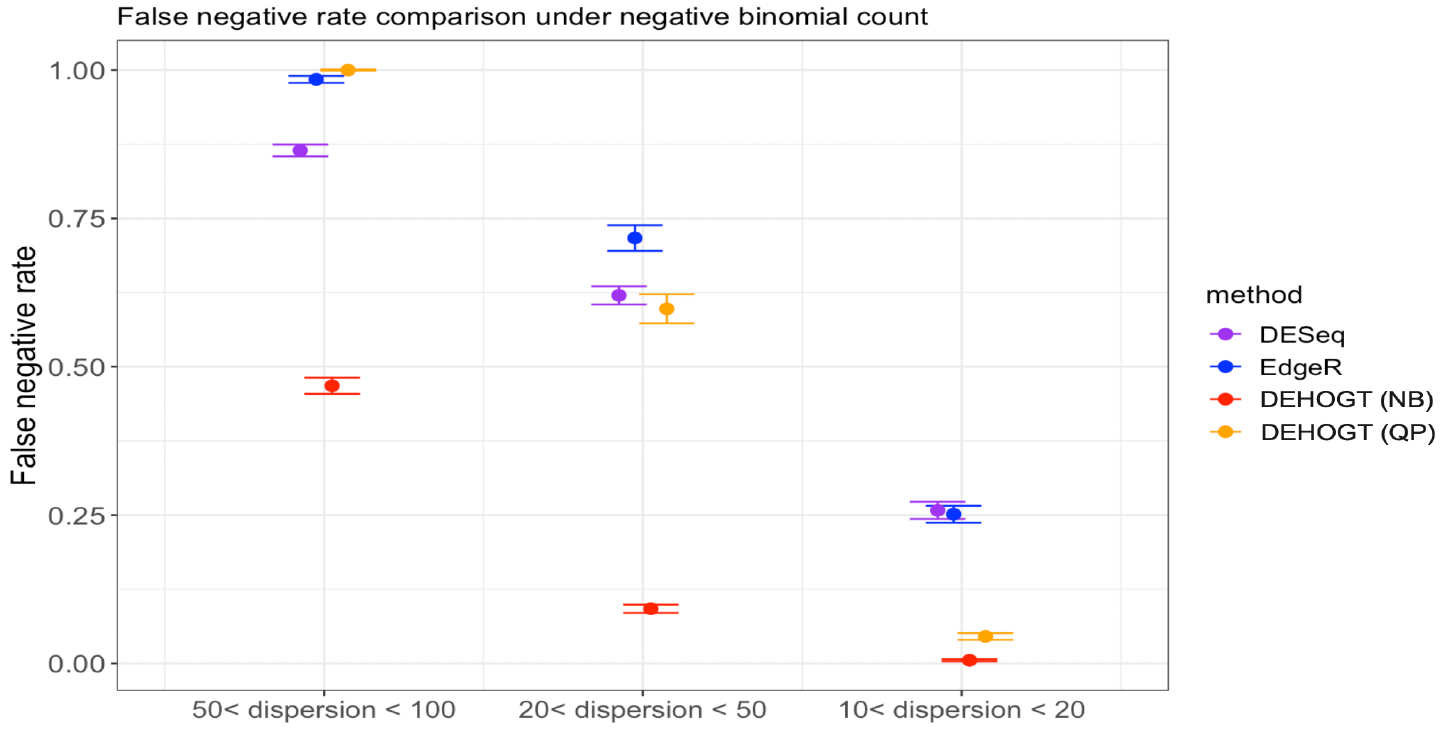
The false negative rate from different methods when the read count follows a negative binomial distribution with different overdispersion levels *θ^NB^*.

The numerical results illustrates that the proposed method DEHOGT has a lower false negative rates than DESeq and EdgeR under different read count generation distributions and different overdispersion levels. Specifically, when the read count distribution is correctly specified, our method consistently achieves lower false negative rate than the EdgeR and DESeq. More importantly, the improvement from the DEHOGT increases as the degree of overdispersion in the read count increases for both quasi-Poisson and negative binominal distributions.

We also investigated the overall discriminative power of the DE gene using different methods when the adjusted p-value cutoff varies between 0 and 1 instead of using 0.05. The overall classification performance is measured by the AUC. The Figures 9 and 8 illustrate the AUC from competing methods under different settings of read counts.

**Figure 8:**
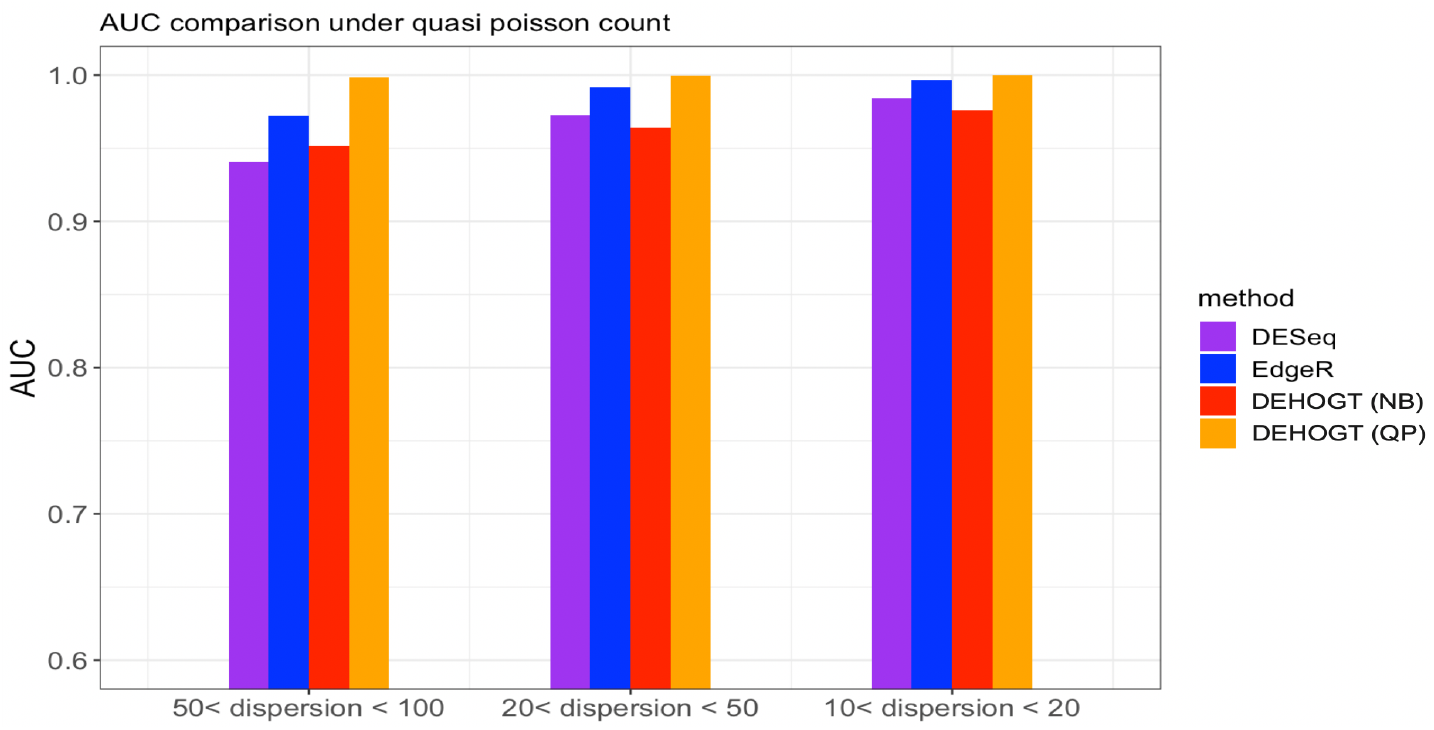
The AUC from different methods when the read count follows the quasi-Poisson distribution with different overdispersion levels *θ^QP^*. The variance of FNR obtained from repeated experiments is illustrated using the bars.

**Figure 9:**
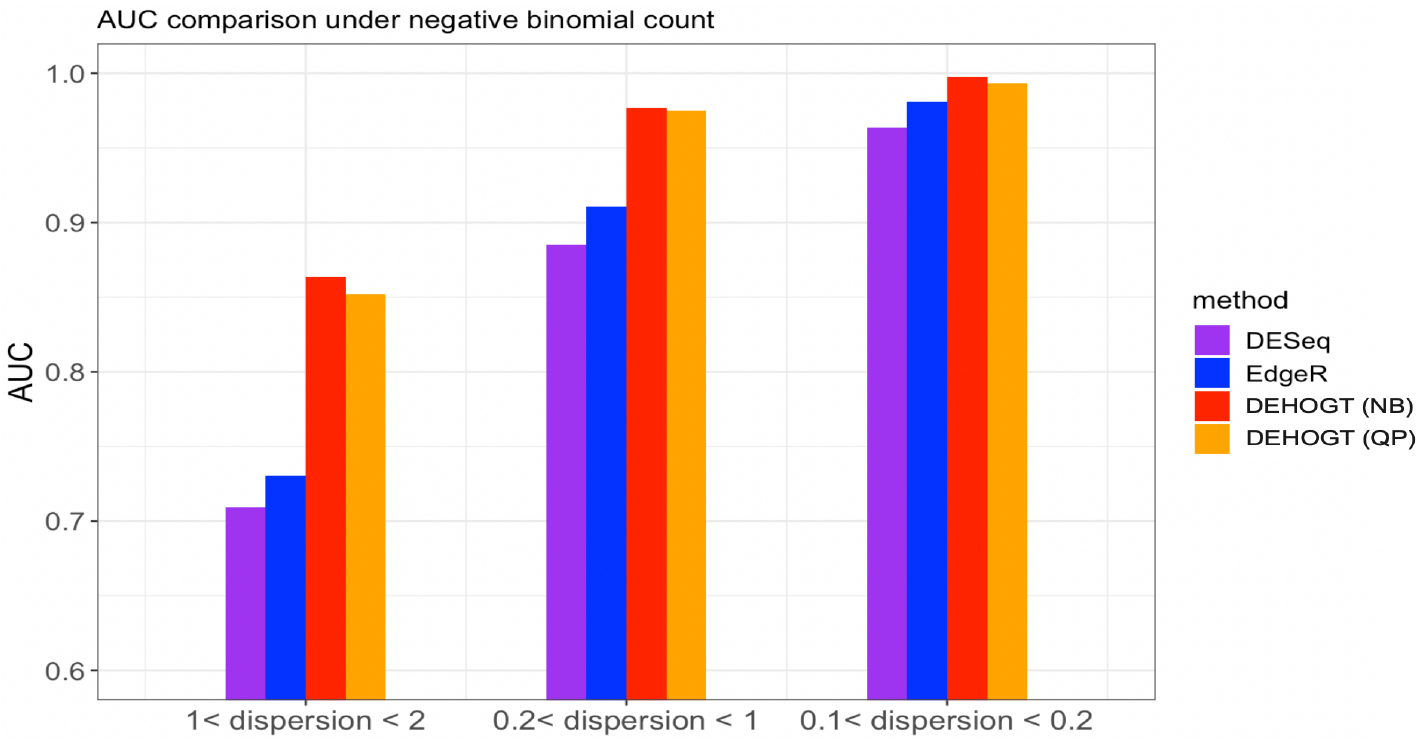
The AUC from different methods when the read counts follow the negative binomial distribution with different overdispersion levels *θ^NB^*.

The above results show that the proposed DEHOGT method achieves a higher AUC in detecting the DE genes than the DESeq and EdgeR, indicating that our method offers a better balance between decreasing false positive rate and false negative rate. In addition, the improvement from our method is more significant as the overdispersion level increases, which is consistent with the aforementioned false negative rate comparison. A higher AUC from the DEHOGT method also implies that it can be more robust against the selection of different cutoff of p-value for DE genes.

We also illustrate the ROC curves in Figure 10 for two representative cases where read counts follow the negative binomial distribution with *θ^NB^* ∈ (1,2), and the quasi-Poisson distribution with *θ^NB^* ∈ (50,100), respectively.

**Figure 10:**
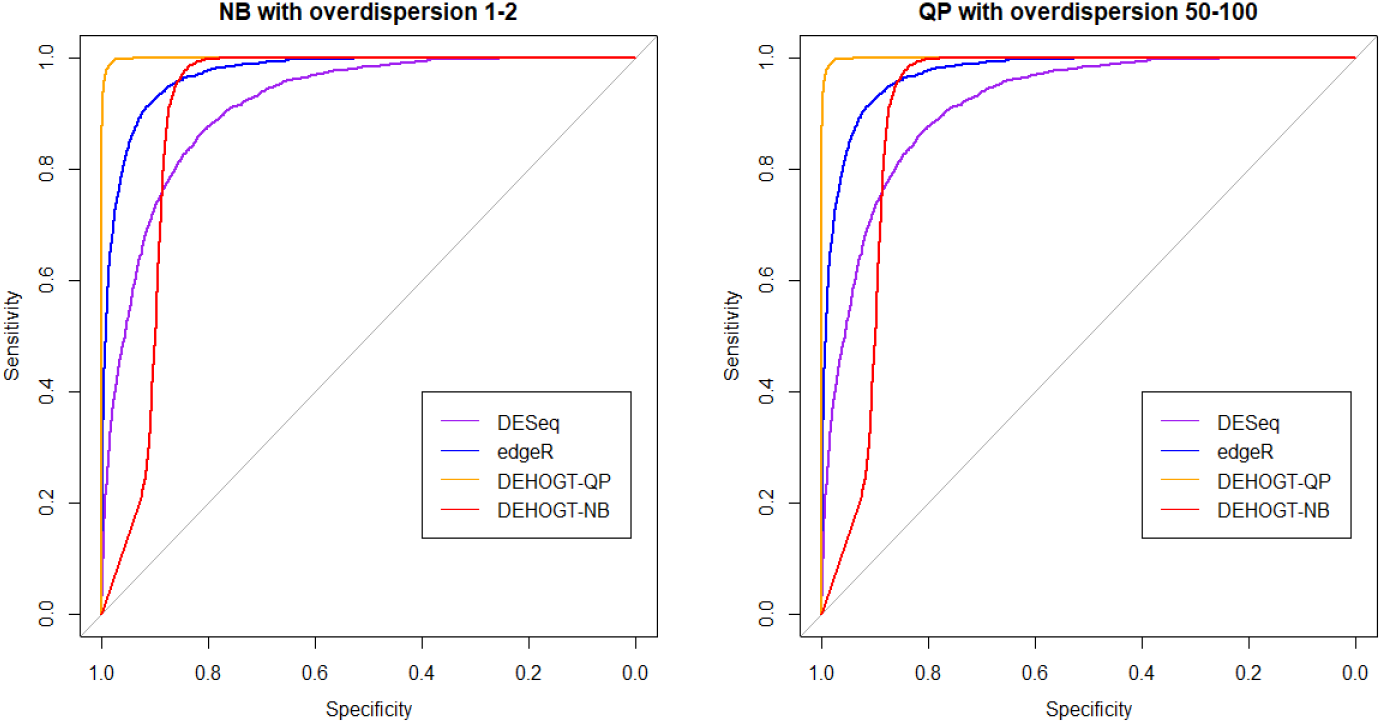
The ROC curve from different methods when the read counts follow the negative binomial distribution with 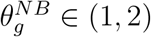 and quasi-Poisson distribution with 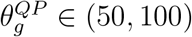.

## 4 Application on Microglia RNA-seq read count data

In this section, we apply the proposed DEHOGT method, DESeq, and EdgeR in the study of post-traumatic stress disorder described in the Introduction section. Specifically, we aim to identify differentially expressed genes from microglia cells that are relevant to the PTSD progress. The RNA-seq data were collected by Uddin research team and Wildman lab at the University of the South Florida. The research performed in-vitro experiments on microglial cells which utilized stress hormones to imitate immune environments similar to PTSD. The function of stress hormones is to adjust the human interior environment, provide energy, and increase heart rate when experience stress [22]. The experiments exposed microglial cells to dexamethasone (dex) and hydrocortisone (cort) serving as stress hormones. The alcohol is also utilized as an additional control treatment to validate if changes in gene expressions are due to the exposure to stress hormones or just a random treatment (alcohol). Specifically, the experiments grew microglial cells under one of the four treatments: hydrocortisone, dexamethasone, alcohol (vehicle), or control. After exposure of three days, RNA-seq data was extracted from the cells on the third day and on the final day of the washout period (day 6), respectively. The goal of study is to identify the genes that are differentially expressed in microglia cells when exposed to different hormones and to determine if the dose of the hormone affects gene expression levels.

More specifically, there are a total of 20,052 expressed genes after quality control preprocessing. There is a total of 9 different treatments with the combination of media (dex, cort, vehicle, and control) and dosage (low and high): dex high, dex low, cort high, cort low, dex vehicle high, dex vehicle low, cort vehicle high, cort vehicle low, and control. On day 3 (time point 3), three repeated samples are collected under each treatment. On day 6 (time point 6), three repeated samples are collected under treatments dex high, dex low, cort high, and cort low, and one sample under dex vehicle high, dex vehicle low, cort vehicle high, and cort vehicle low.

We first investigated the level of empirical dispersion in the microglia RNA-seq read counts. Specifically, we examine the relation between sample count mean and sample count variance across all genes. Figure 11 illustrates a quadratic growth of count variance over count mean. In addition, we fit a quadratic regression on count variance over count mean, where an adjusted *R*^2^ coefficient reaches 0.66. Therefore, we choose to use a negative binomial distribution as the read counts generating process in the proposed DEHOGT method.

**Figure 11:**
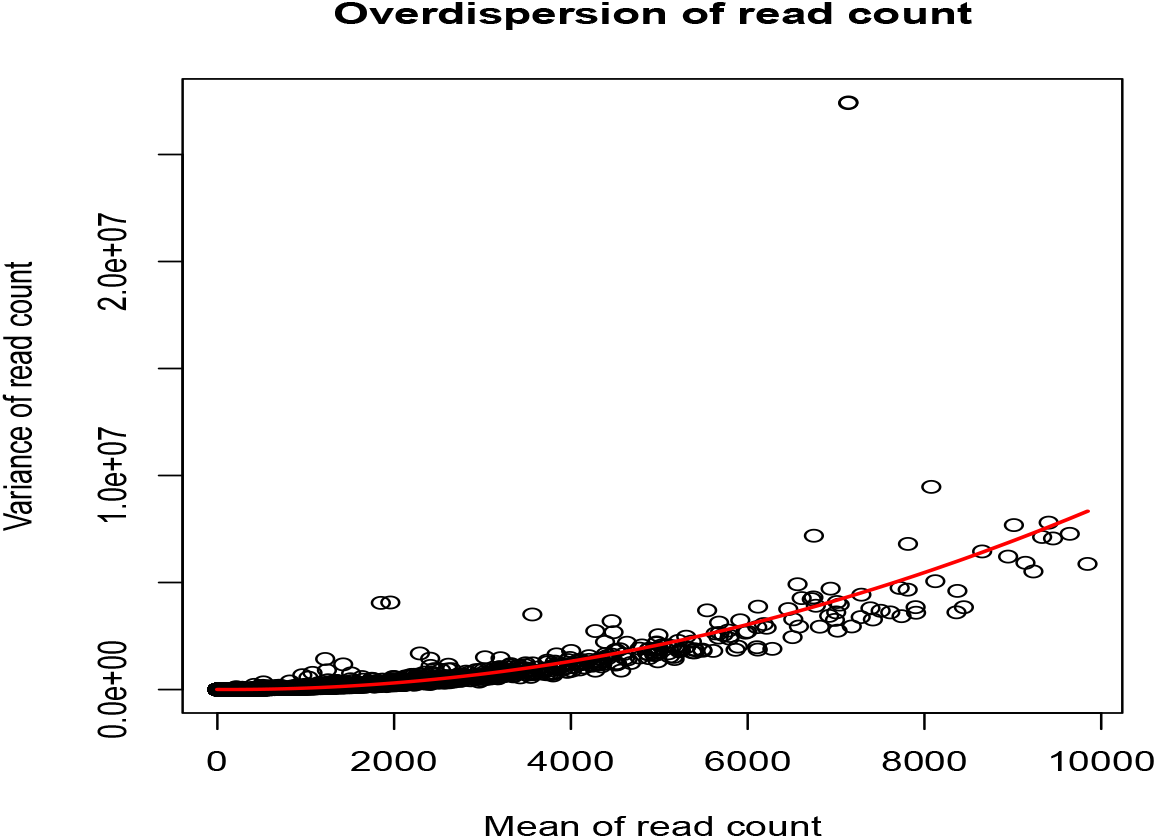
The dispersion of genewise RNA read counts. Each dot corresponds to a sample count from a specific gene.

We utilize DEHOGT, DESeq, and EdgeR to select DE genes under the following 7 treatment comparison pairs: dex high at time point 3 and control (dexh3 vs control), dex high at time point 6 and control (dexh6 vs control), cort high at time point 3 and control (corth3 vs control), cort high at time point 6 and control (corth6 vs control), dex vehicle high and dex high at time point 3 (dexvh3 vs dexh3), dex vehicle low and dex low at time point 3 (dexvl3 vs dexl3), cort vehicle high and cort high at time point 3 (cortvh3 vs corth3). In selecting DE genes between the two treatments, we follow the criterion in Section 2 in that the adjusted p value is smaller than 0.05, and the log2fold change is larger than 1.5.

We first illustrate the number of DE genes selected by competing methods. Table 1 shows that the proposed method tends to select more genes than the other two methods, especially compared to the DESeq. In the exploratory stage, it is critical to include as many relevant genes as possible for the downstream analysis. The DEHOGT method is more effective in reducing the false negative rate in detecting PTSD-related genes by identifying a larger candidate pool of DE genes.

**Table 1:** The number of selected DE genes from microglia RNA-seq data under different treatment comparisons.

| Methods | Treatment Pairs |  |  |  |  |  |  |
| --- | --- | --- | --- | --- | --- | --- | --- |
|  | dexh3 vs control | dexh6 vs control | corth3 vs control | corth6 vs control | dexvh3 vs dexh3 | dexvl3 vs dexl3 | cortvh3 vs corth3 |
| DESeq | 221 | 45 | 166 | 1 | 255 | 118 | 112 |
| EdgeR | 419 | 115 | 308 | 3 | 469 | 180 | 216 |
| DEHOGT | 981 | 237 | 355 | 582 | 383 | 148 | 256 |

We conduct detailed analysis for the DE genes based on three methods for each treatment pair. In general, we investigate the overlapping in DE genes from three methods, where the findings are illustrated via the Venn diagram in Figure 12 to Figure 18. Notice that the proposed DEHOGT method selects more DE genes than DESeq and EdgeR for all pairwise comparisons between treatments except dexvh 3 vs dexh 3 and dexvl 3 vs dexl 3, demonstrating that the proposed method can identify more DE genes to reduce the potential risk of missing underlying relevant genes. In the following, we provide an interpretation for the treatment pair dexh6 versus control. The interpretation of other treatment pairs can be conducted similarly. The Venn diagram in Figure 13 shows that all the DE genes selected by the DESeq are also selected by EdgeR, and 86.7% of the DE genes selected by DESeq are also selected by DEHOGT. In addition, 61.7% of the DE genes selected by EdgeR are detected by DEHOGT.

**Figure 12:**
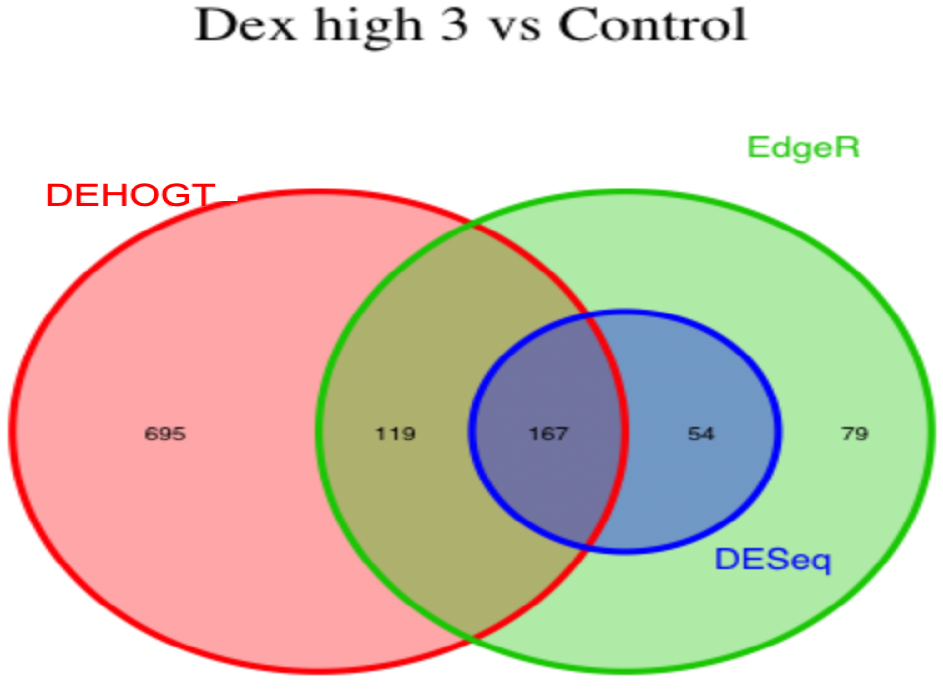
The selected DE genes from DEHOGT, DESeq, EdgeR under treatment comparison dexh3 versus control.

**Figure 13:**
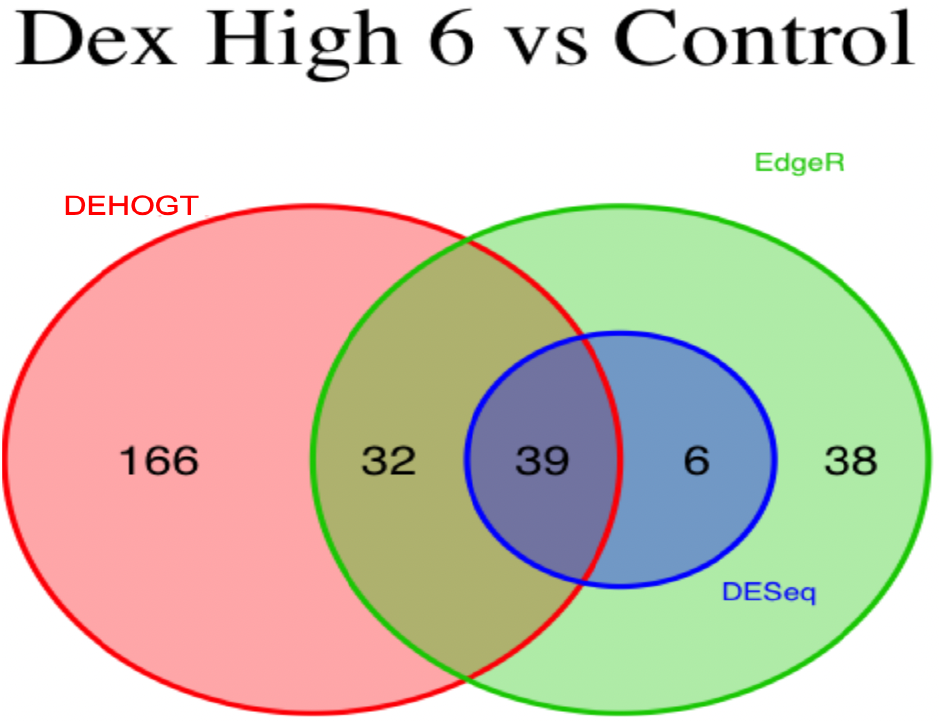
The selected DE genes from DEHOGT, DESeq, EdgeR under treatment comparison dexh6 versus control.

**Figure 14:**
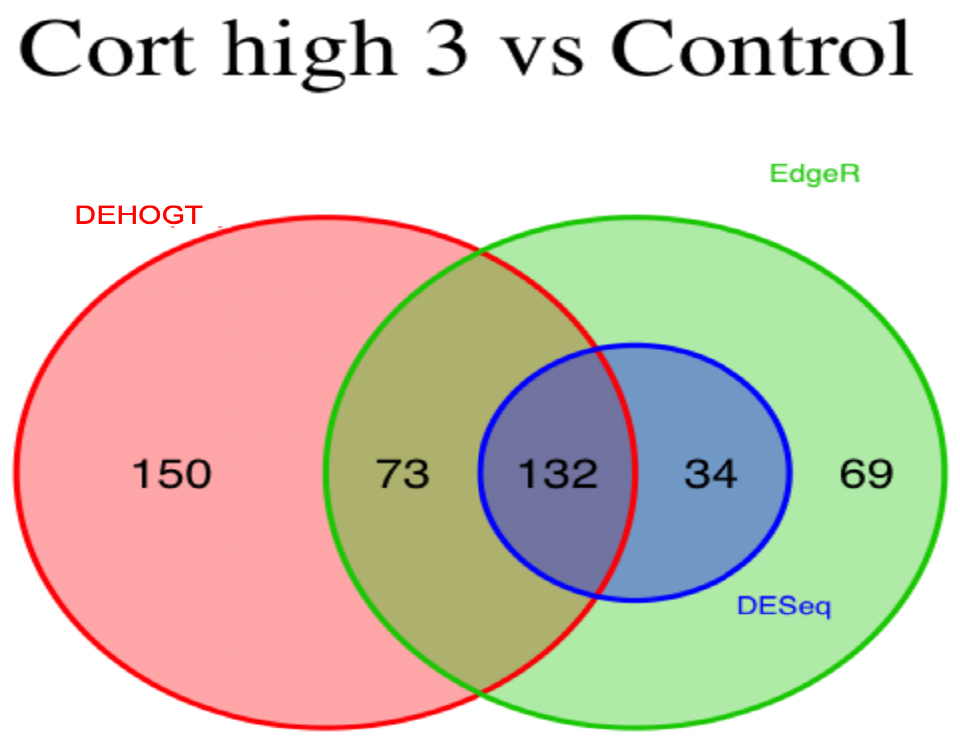
The selected DE genes from DEHOGT, DESeq, EdgeR under treatment comparison corth3 versus control.

**Figure 15:**
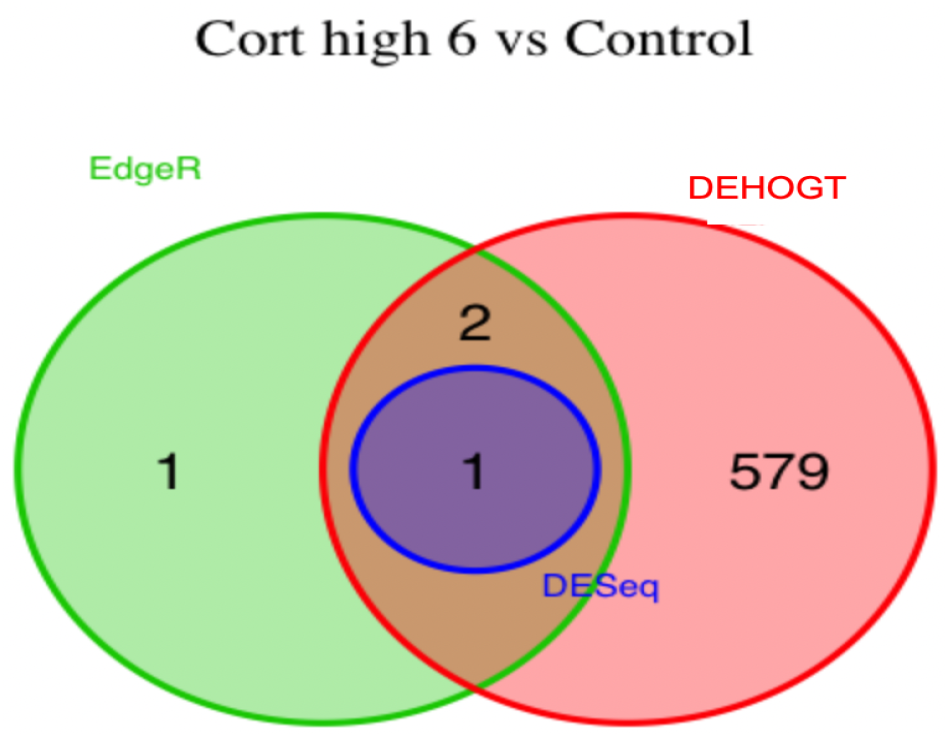
The selected DE genes from DEHOGT, DESeq, EdgeR under treatment comparison corth6 versus control.

**Figure 16:**
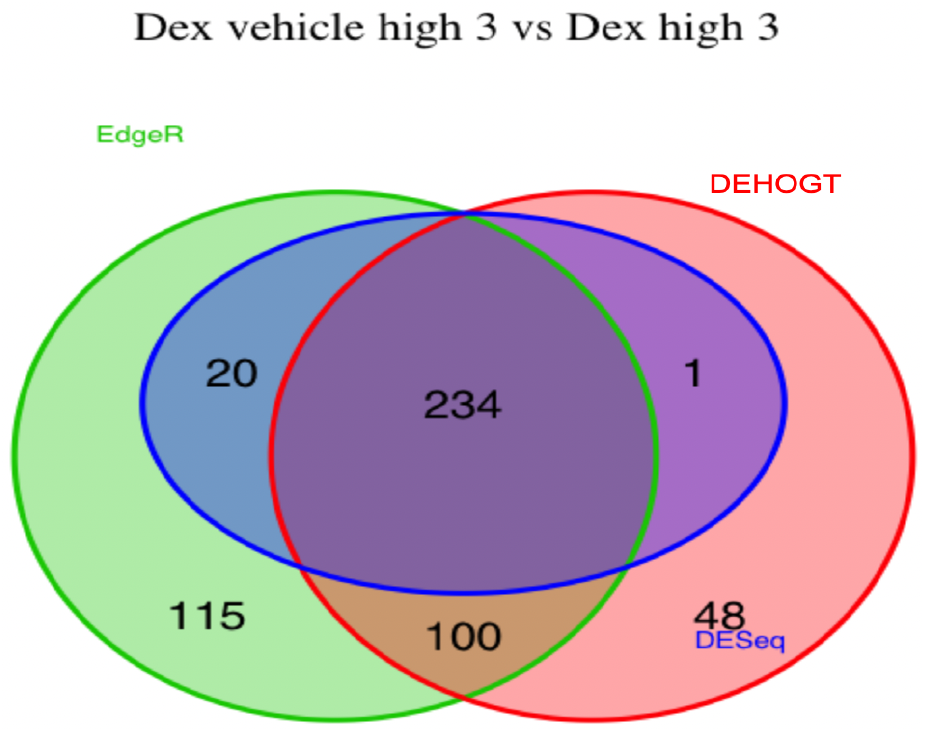
The selected DE genes from DEHOGT, DESeq, EdgeR under treatment comparison dexvh3 versus dexh3.

**Figure 17:**
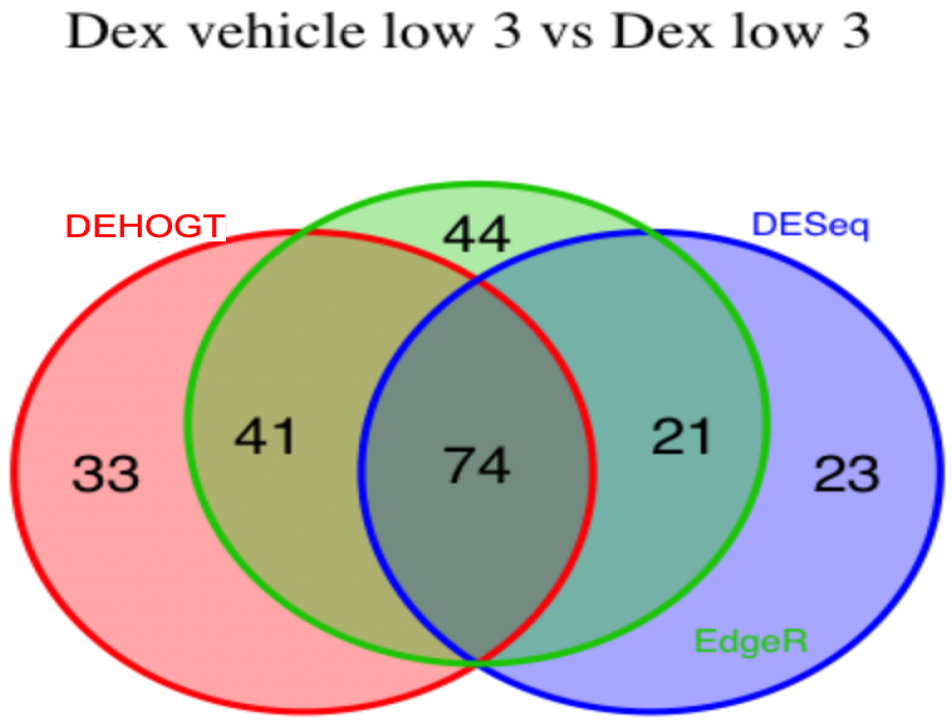
The selected DE genes from DEHOGT, DESeq, EdgeR under treatment comparison dexvl3 vs dexl3.

**Figure 18:**
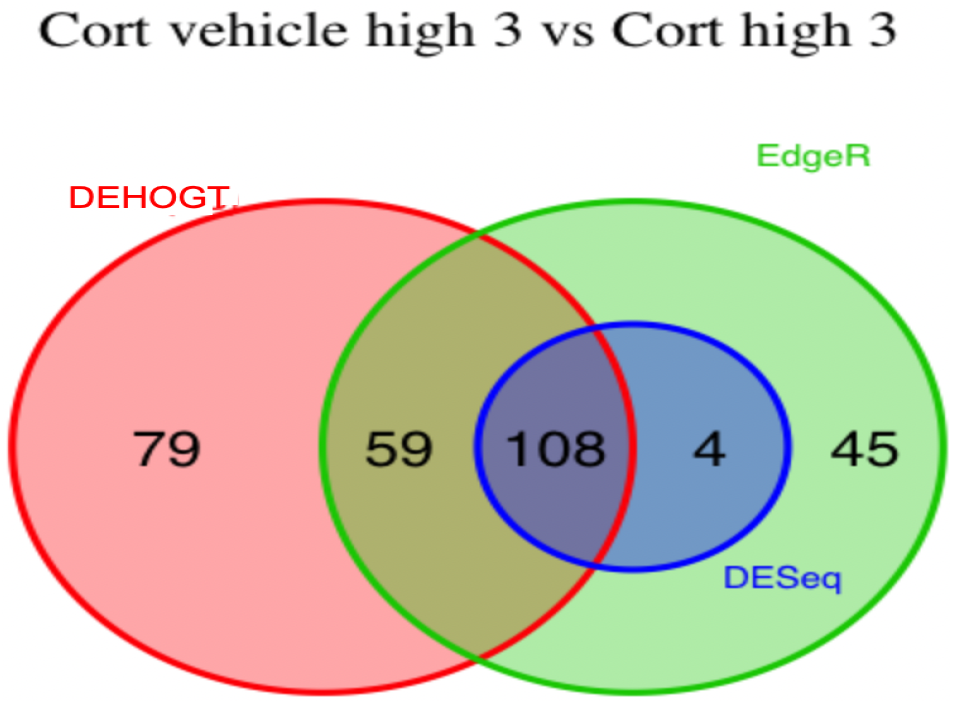
The selected DE genes from DEHOGT, DESeq, EdgeR under treatment comparison cortvh3 vs corth3.

The proposed method identifies three genes *CRISPLD2, TSC22D3*, and *PSG1* which are differentially expressed under the three treatment comparisons: dexvh 3 versus dexh 3, dexvl 3 versus dexl 3, and cortvh3 versus corth3. Specifically, the glucocorticoid-responsive gene *CRISPLD2* is found to be differentially expressed in read counts from an RNA-seq experiment with muscle cells exposed to dexamethasone [10]. The another glucocorticoid-responsive gene *TSC22D3 (GILZ*) is found to be differentially expressed under gonorrhea or chlamydia exposure based on many animal and human gene studies that examine different cell types [7, 9]. These evidences support the fact that *TSC22D3* serves as a mediator for the anti-inflammatory activity of gonorrhea or chlamydia summarized in [26]. The gene *PSG1* is found to activate the underlying beta 1 (TGF-β1) known as transforming growth factor, which is an essential cytokine process in suppression and immunoregulation of inflammatory T cells [28, 5].

We also list the significant DE genes uniquely selected by the three methods in Table 2, which demonstrates that most of the DE genes identified by DESeq are also selected by DEHOGT and EdgeR. Specifically, the gene *FKBP5* is identified by the proposed method but not identified by the other methods under the comparison dexh3 versus dexvh3. The gene *FKBP5* is a co-chaperone adjust the activity of glucocorticoid receptor. *FKBP5* is known as an important modulator of responding stress. In many studies using different cell types, the dysregulation phenomenon of *FKPB5* is found in many stress-related psychopathologies via investigating single nucleotide polymorphisms [4, 2], gene expression [11], and DNA methylation profiles [14].

**Table 2:**
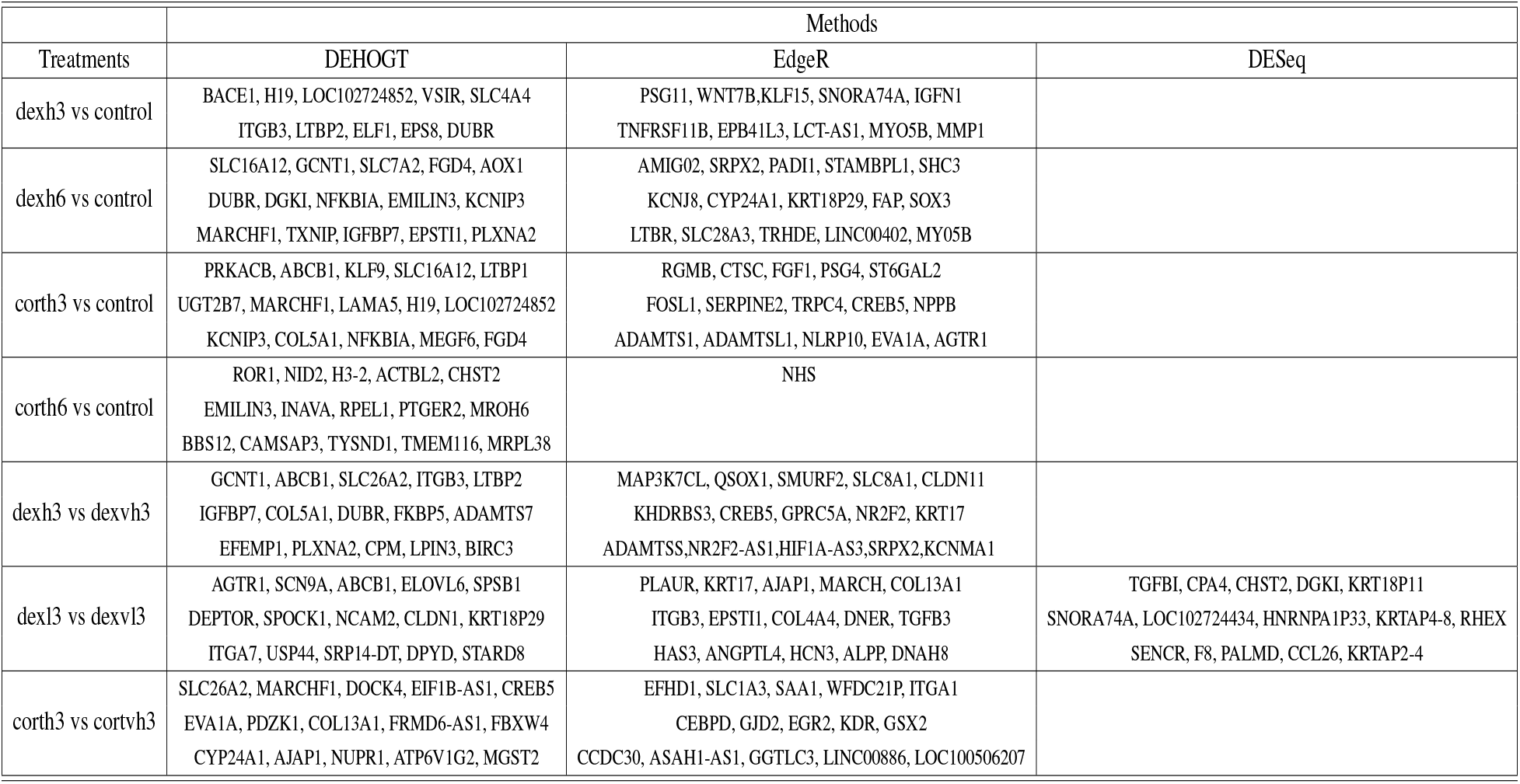
Top 15 unique DE genes unique selected by DEHOGT, EdgeR, and DESeq.

In addition, we examine the most significant DE genes among the overlaps of the three methods in Figure 19 to Figure 24. For treatment pair dexh6 versus control, Figure 20 lists the 30 most significant DE genes which are overlapping for all three methods, and the bar charts with different colors represent the rank of p-values from the three methods. A shorter bar indicates a smaller p-value and therefore a more significantly differentially expressed genes under dex high and control comparison. The DEHOGT method selects genes *ROR1, FAT3, TLR4, CERNA2, ADPRHL1, NID2, CRISPLD2*, and *ABCA8* as the top 8 significant DE genes, and these genes are also among the top significant DE genes selected by EdgeR and DESeq. In general, our method provides a list of the top significant DE genes which is consistent with the DESeq and EdgeR in comparing dexh6 versus control. This cross-validation on DE genes via the three methods confirms the association between PTSD and the top DE genes which are identified by the DEHOGT. In particular, the previously mentioned genes *TSC22D3* and *PSG1* are identified by all three methods for the vehicle treatment comparisons: dexvh 3 versus dexh 3, dexvl 3 versus dexl 3, and cortvh3 versus corth3. These results provide evidence of further need to explore their roles in the formulation of PTSD.

**Figure 19:**
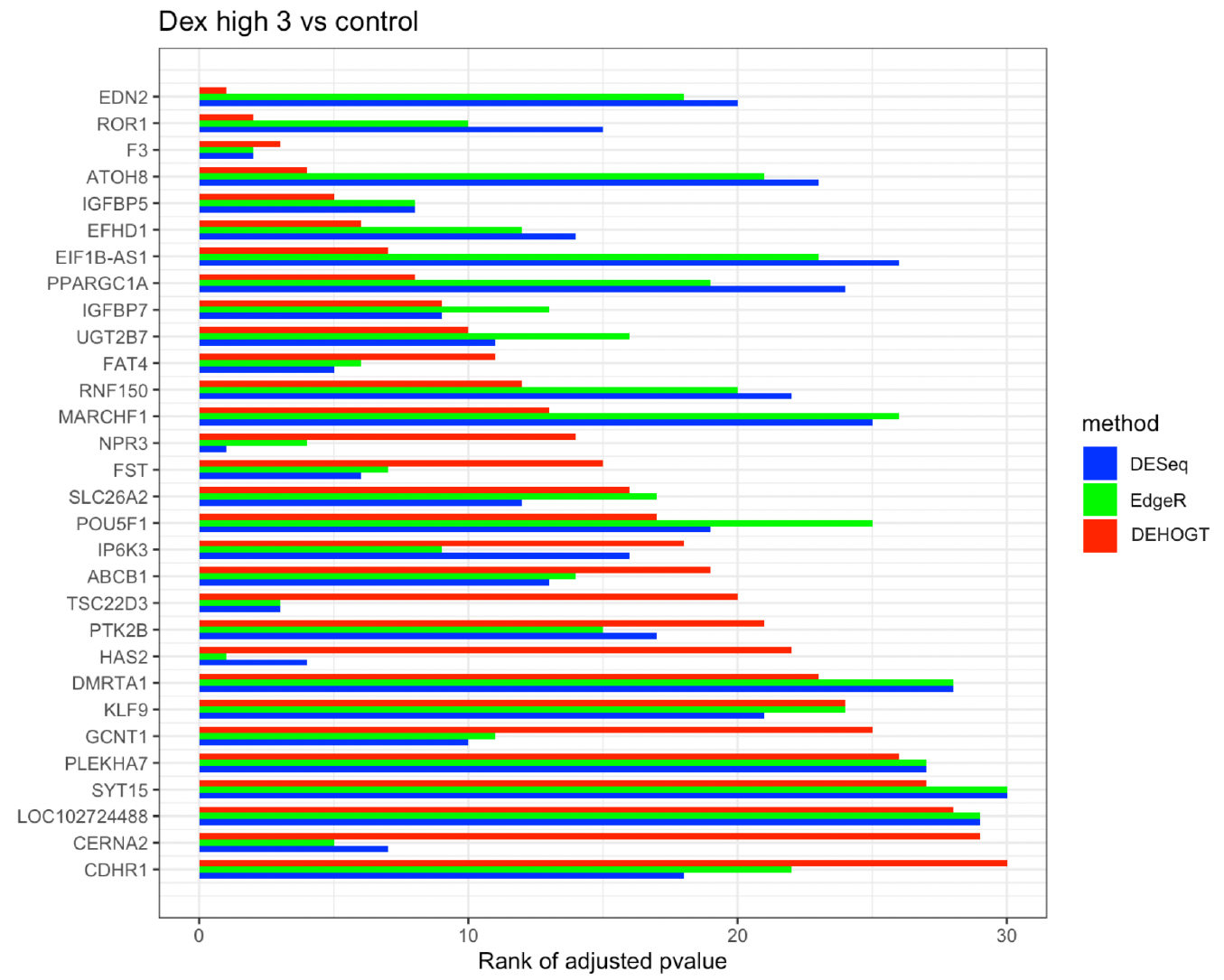
The rank of p-value of selected genes under treatment comparison dexh3 versus control, a shorter bar indicates a smaller p-value (more significantly differently expressed).

**Figure 20:**
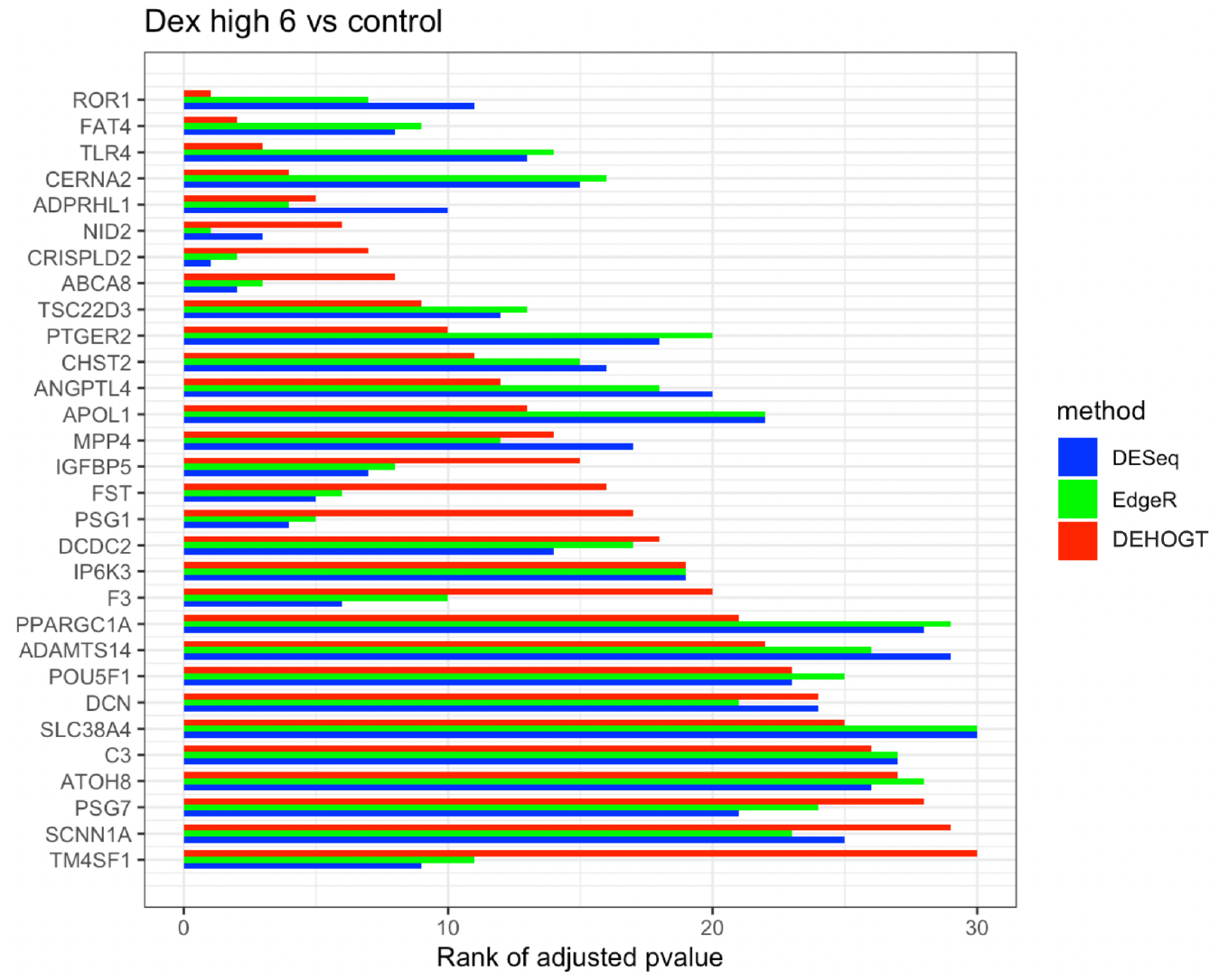
The rank of p-value of selected genes under treatment comparison dexh6 versus control, and shorter bar indicates a smaller p-value.

**Figure 21:**
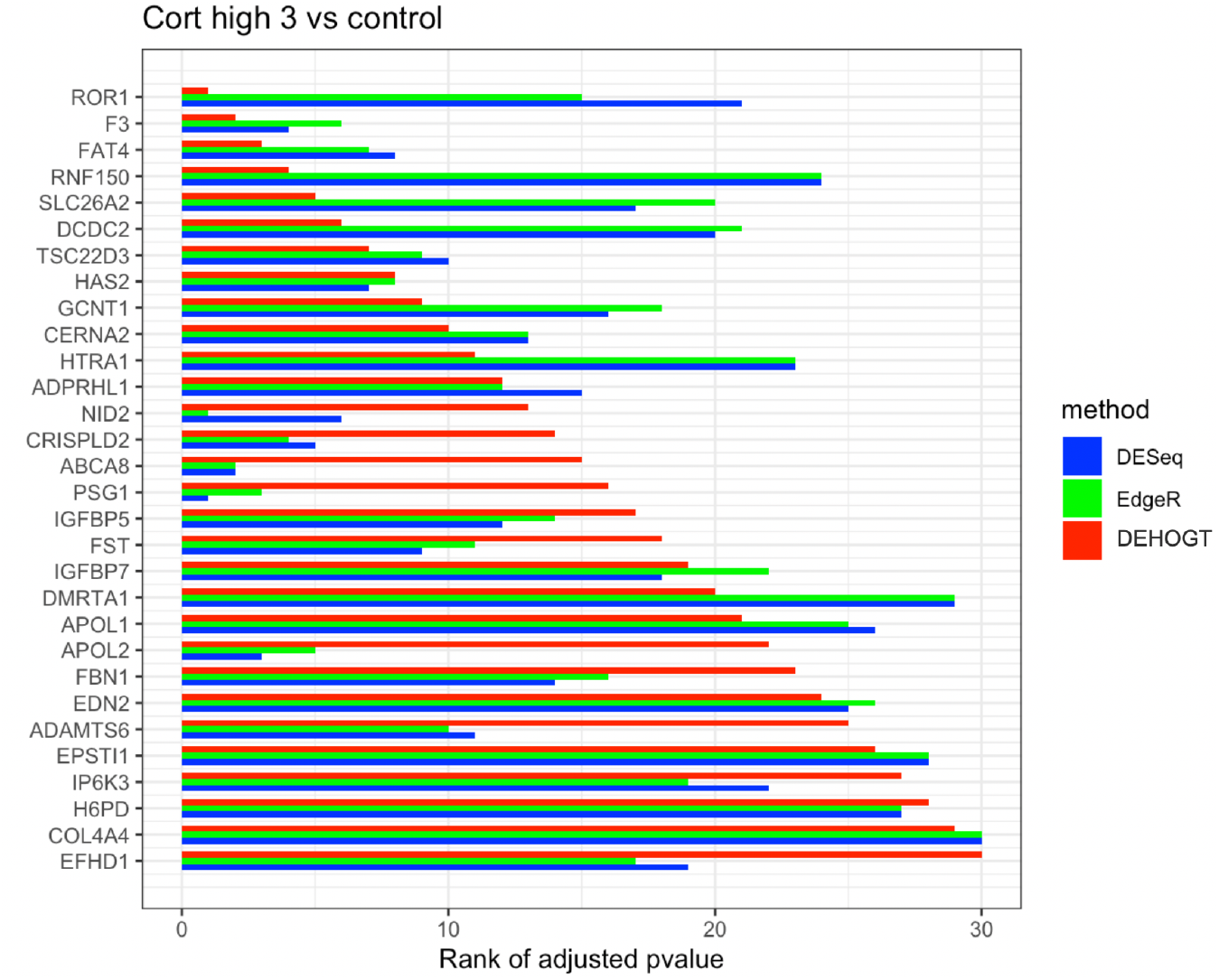
The rank of p-value of selected genes under treatment comparison corth3 versus control, and shorter bar indicates a smaller p-value.

**Figure 22:**
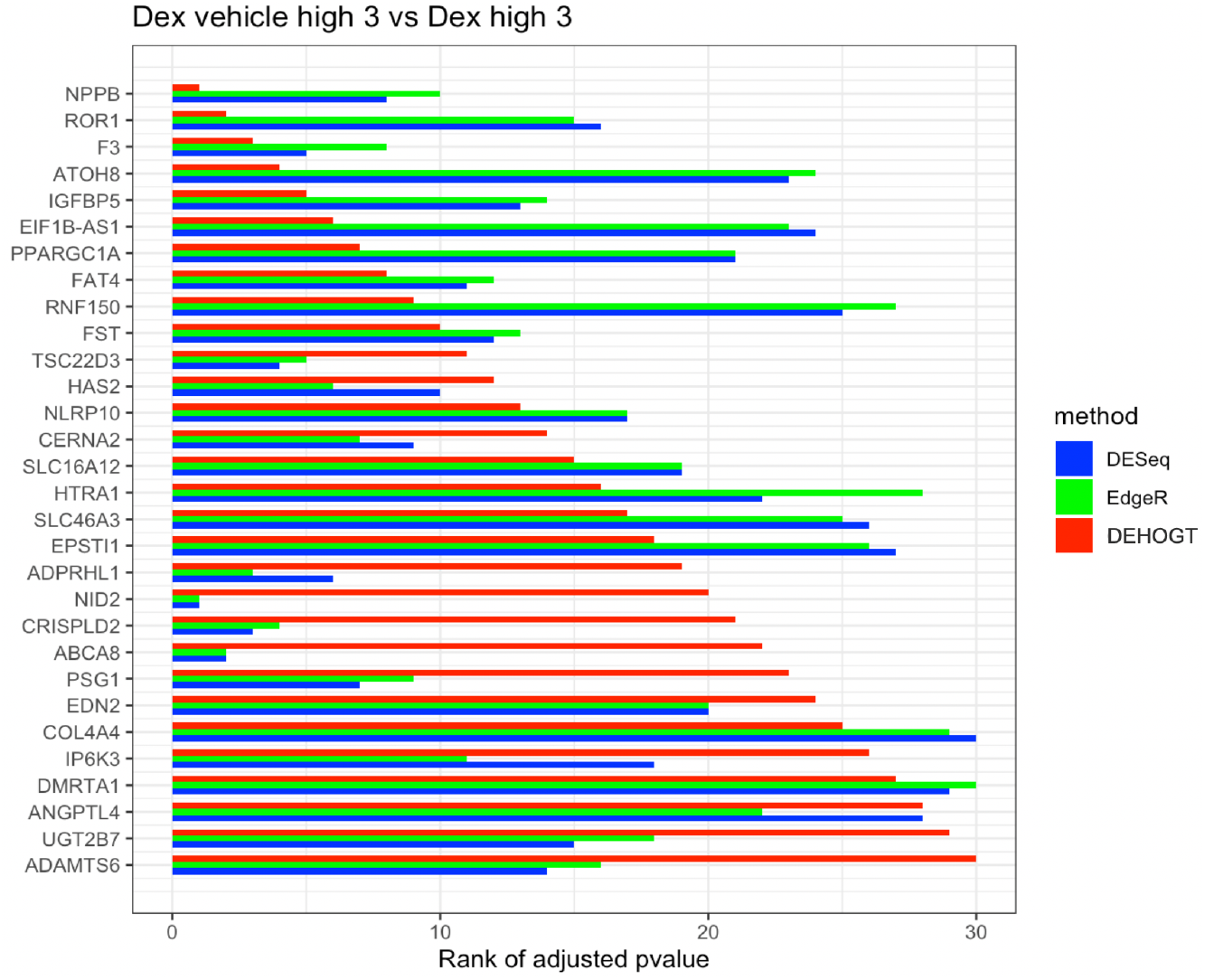
The rank of p-value of selected genes under treatment comparison dexvh versus dexh, and shorter bar indicates a smaller p-value.

**Figure 23:**
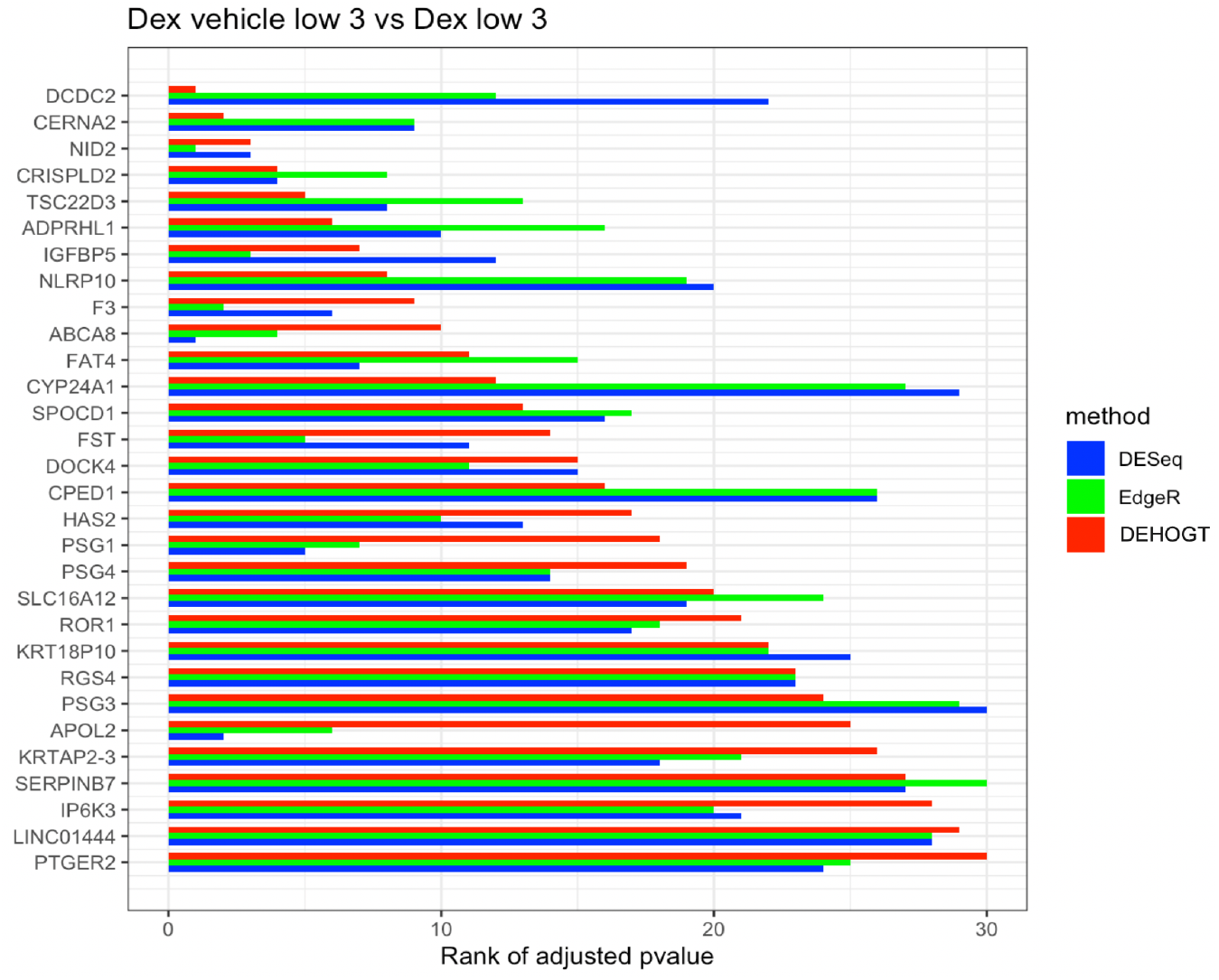
The rank of p-value of selected genes under treatment comparison dexvl versus dexl, and shorter bar indicates a smaller p-value.

**Figure 24:**
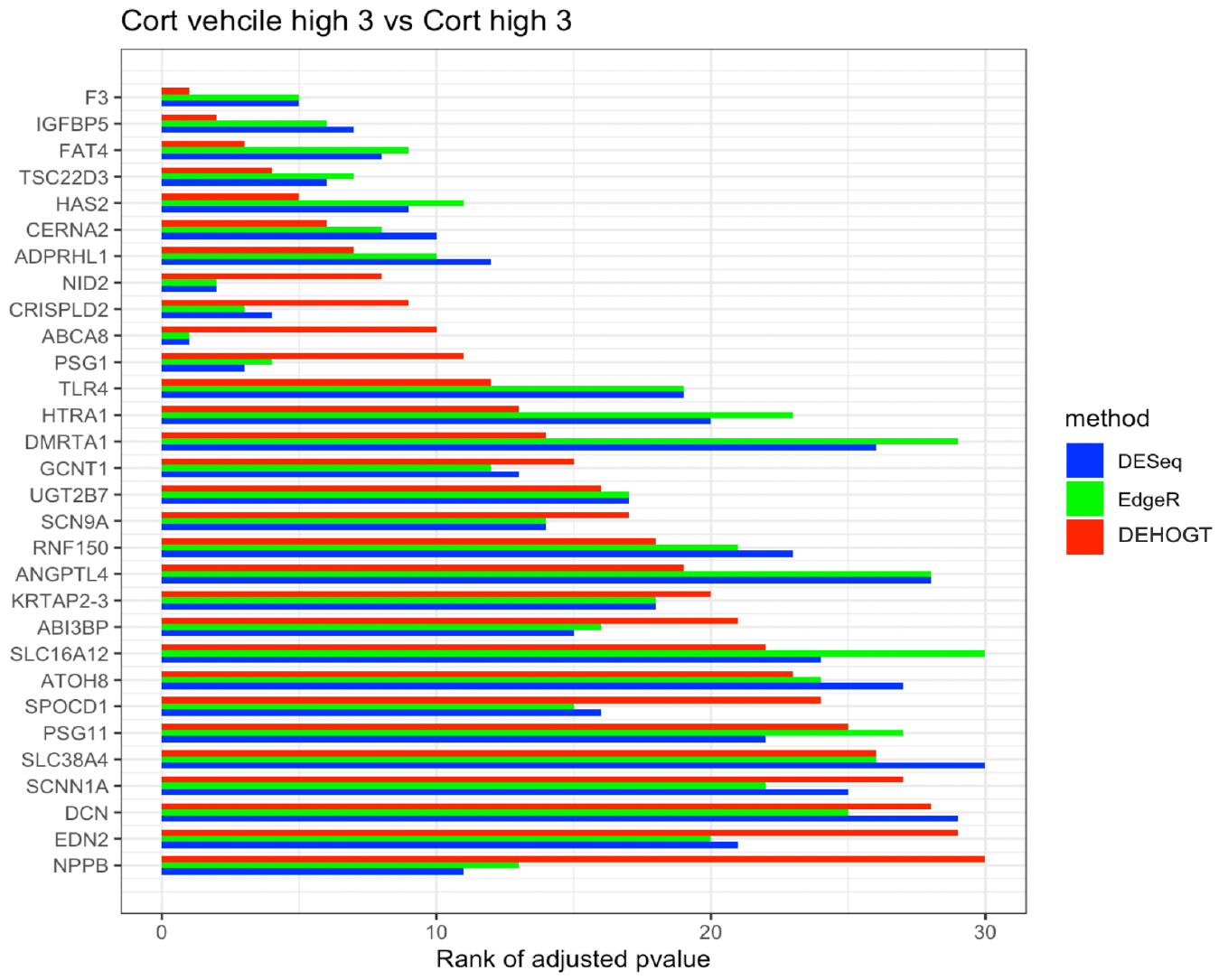
The rank of p-value of selected genes under treatment comparison cortvh versus corth, and shorter bar indicates a smaller p-value.

## 5 Discussion

In this paper, we propose a revised differential expression analysis procedure DEHOGT for identifying differentially expressed genes based on overdispersed RNA-seq read count data. DEHOGT adopts a joint estimation of logfold changes that incorporates samples from all treatments simultaneously to utilize cross-treatment information. In addition, the proposed method takes advantage of within-treatment independence structures among genes to increase the effective sample size, which leads to stronger power in detecting DE genes. Furthermore, our method enjoys flexibility in utilizing different read count generating distributions instead of fixing only one negative binominal distribution as in the popular methods such as EdgeR and DESeq. This allows us to choose a generating distribution adopted to the empirical dispersion level. Therefore, DEHOGT has the potential to be applied for other genetic datasets with similar challenges of heterogeneous overdispersion levels.

In our simulation study, we demonstrate that the proposed method achieves better performance in detecting DE genes compared with the EdgeR and DESeq methods, especially when the per-treatment sample size is relatively small. The numerical experiments suggests that DEHOGT is less conservative in selecting DE genes due to adopting the individual fitting procedure. This property enables our method to have improved performance in controlling the false negative rate which is more critical for downstream analysis.

We further apply our method and compare it with EdgeR and DESeq on a real application in a microglia RNA-seq dataset collected by our team. Specifically, our method identifies more potential genes which may be potentially more relevant to PTSD than either EdgeR and DESeq.

In addition, the cross-validation among EdgeR, DESeq and the proposed method provides a rich and robust candidate pool for genes relevant to PTSD. These results were obtained in the microglia dataset despite having issues of overdispersion and small sample size.

The popular existing methods DESeq and EdgeR identify differentially expressed genes by adopting an aggregate estimation strategy for read count overdispersion levels, which relies on the key assumption that genes with similar expression levels have similar overdispersion levels. The numerical results in this paper indicate that this assumption might be questionable under the scenario when heterogeneity of gene expression level is high. The violation of this assumption can undermine the detection power of methods based on aggregate estimators of overdispersion, especially when the overdispersion level is high. In contrast, estimating overdispersion levels for each gene separately can be more robust under high heterogeneity in gene expressions. On the other hand, the proposed independent estimation scheme integrates samples from different treatments instead from different genes, which might lose a certain amount of statistical testing power especially when the sample size is small. One direction worth of further exploration is to incorporate neighborhood similarity structures among genes such that the overdispersion estimation of a spe-cific gene can borrow the information of samples from correlated genes, therefore we can increase the effective sample size for estimating overdispersion levels. A potential strategy could utilize gene-wise covariate variables or develop an adaptive fused-type penalty on gene overdispersion levels.

## 6 Appendix: Microglia cell experiment design

The microglial cell line HMC3 (ATCC CRL-3304, Manassas, Virginia) was used for in vitro experimentation following successful cell line authentication and Mycoplasma testing (Genetica, Burlington, NC). HMC3 cells (passage eight) were seeded in T-25 flasks with 2 x 105 viable cells and incubated at 37°C and 5% CO2. After 24 hours, the growth medium in each T-25 was replaced with one of the following treatments: dexamethasone (1 or 0.01 μM), hydrocortisone (10 or 0.01 μM), vehicle (ethanol alcohol) or control (untreated media). Cells were incubated in treatment media for three days at 37°C and 5% CO2 and imaged daily using the Axio Vert.A1 inverted microscope (Zeiss Oberkochen, Germany). At three days post-exposure (D3), cells were collected from each flask individually, quantified on the Countess II cell counter (Invitrogen Waltham, MA), seeded at 2 x 105 viable cells/flask in new T-25 flasks with normal growth medium, and incubated for three additional days (i.e., washout period). The remaining D3 cell suspension for each flask was divided equally between two microcentrifuge tubes, pelleted and washed with PBS. One cell pellet per flask was placed in −80°C storage for future DNA extraction; the remaining cell pellet underwent RNA extraction using the RNeasy Mini Kit (QIAGEN, Hilden, Germany) protocol adapted for the QIAcube automated system (QIAGEN). On the final day of the washout period (D6), cells from each flask were imaged, collected in suspension and then quantified on the Countess II. Cell suspensions were split equally into two aliquots and then prepped for nucleic acid extraction as described for D3.

RNA samples from D3 and D6 were DNase treated (Dnase I kit; Sigma), quantified on the Qubit (RNA BR Assay Kit; Invitrogen) and scored for RNA integrity on the TapeStation (High Sensitivity RNA ScreenTape; Agilent). Library preparation was performed following the Illumina TruSeq Stranded Total RNA Library Prep Kit protocol (Illumina, San Diego, CA) with TruSeq RNA Single Indexes (Set A and B; Illumina). Library quantity and quality were assessed using the Qubit 1X dsDNA HS Assay (Invitrogen), TapeStation High Sensitivity D1000 ScreenTape (Agilent), and using the KAPA Library Quantification Kit (Roche Basel, Switzerland) for the LightCycler 96 (Roche). RNA sequencing was conducted on the NextSeq 550 (Illumina) using the High Output Kit with 76 paired-end cycles (Illumina).

## 7 Acknowledgements

The authors would like to acknowledge the USF Genomics Core for their support of the microglia cell experiment.

## 8 Availability of data and materials

The datasets analysed during the current study are available in the NCBI’s Gene Expression Omnibus (GEO) repository and are accessible through GEO Series accession number GSE219208 and link https://www.ncbi.nlm.nih.gov/geo/query/acc.cgi?acc=GSE219208.

The experiment description and algorithm implementation are available via the following weblinks: https://github.com/xiaobai0518/DEHOGT. Operating systems: Windows, Linux, MacOS Programming language: R. Other requirements: RStudio. License: GPL-3.0.

